# CRISPR-Cas12a REC2 – NUC interactions drive target-strand cleavage and constrain trans cleavage

**DOI:** 10.1101/2025.03.23.644851

**Authors:** Anthony Newman, Aakash Saha, Lora Starrs, Pablo R. Arantes, Giulia Palermo, Gaetan Burgio

## Abstract

CRISPR-Cas12a effects RNA-guided cleavage of dsDNA in *cis*, after which it remains catalytically active and non-specifically cleaves ssDNA in *trans*. Native host-defence by Cas12a employs *cis* cleavage, which can be repurposed for the genome editing of other organisms, and *trans* cleavage can be used for *in vitro* DNA detection. Cas12a orthologues have high structural similarity and a conserved mechanism of DNA cleavage, yet highly different efficacies when applied for genome editing or DNA detection. By comparing three well characterised Cas12a orthologues (FnCas12a, LbCas12a, and AsCas12a), we sought to determine what drives their different *cis* and *trans* cleavage, and how this relates to their applied function.

We integrated *in vitro* DNA cleavage kinetics with molecular dynamics simulations, plasmid interference in *E. coli*, and genome editing in human cell lines. We report large differences in *cis* cleavage kinetics between orthologues, which may be driven by dynamic REC2-NUC interactions. We generated and tested REC2 and NUC mutants, including a hitherto unstudied ‘NUC loop’, integrity of which is critical for the function of Cas12 orthologues. In total, our *in vitro, in vivo,* and *in silico* survey of Cas12a orthologues highlights key properties that drive their function in biotechnology applications.

**Graphical abstract:** 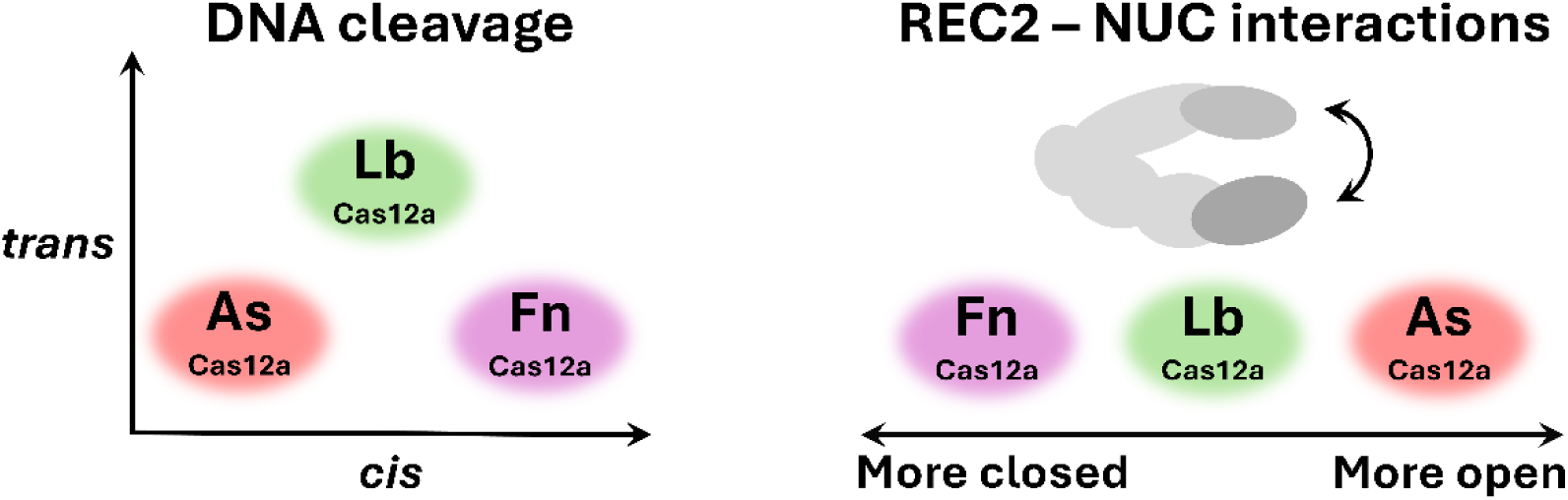

## Introduction

CRISPR-Cas (<u>c</u>lusters of regularly interspaced <u>s</u>hort <u>p</u>alindromic repeats-<u>C</u>RISPR-<u>as</u>sociated) are adaptive immune systems in bacteria and archaea that interfere with foreign nucleic acid sequences in an RNA-guided fashion (1). Cas12a, formerly named Cpf1, is the signature effector of type V-A CRISPR systems (2). In host defence, Cas12a binds to a guide RNA derived from its CRISPR array – the crRNA – to effect RNA-programmable cleavage of double-stranded DNA in *cis* (3–5). This RNA-guided nuclease activity has been widely employed for the genome editing of eukaryotic cells (5–7). *In vitro*, Cas12a remains catalytically active after *cis* cleavage, and can cut ssDNA, RNA, and nick dsDNA (8–10). The target-activated *trans* cleavage of Cas12a underlies its applications for molecular detection. With reverse transcription and aptamer strategies, RNA, proteins, small molecules, and even heavy metals can also be detected using Cas12a (11, 12).

Cas12a assumes a ‘crab-claw’ structure of two lobes; with nuclease (NUC) and recognition (REC) lobes that effect their eponymous functions (4, 13–18) (**Figure 1A**). Cas12a scans double-stranded DNA and initiates R-loop formation at protospacer-adjacent motifs (PAMs) (19). The REC lobe recognises a matching DNA target site by stable R-loop formation between crRNA and a hybridised DNA strand (target-strand, TS) (14, 18, 20). Docking of the flexible REC domain to the bridge helix (BH) domain activates the distant RuvC domain in the NUC lobe, by stabilising the open conformation of a RuvC-occluding ‘lid’ loop (18, 20–24). This narrow active site cleft can only sterically accommodate single-stranded DNA, so dsDNA cleavage by Cas12a has to occur via sequential cleavage of unwound strands (4, 13, 25).

**Figure 1:**
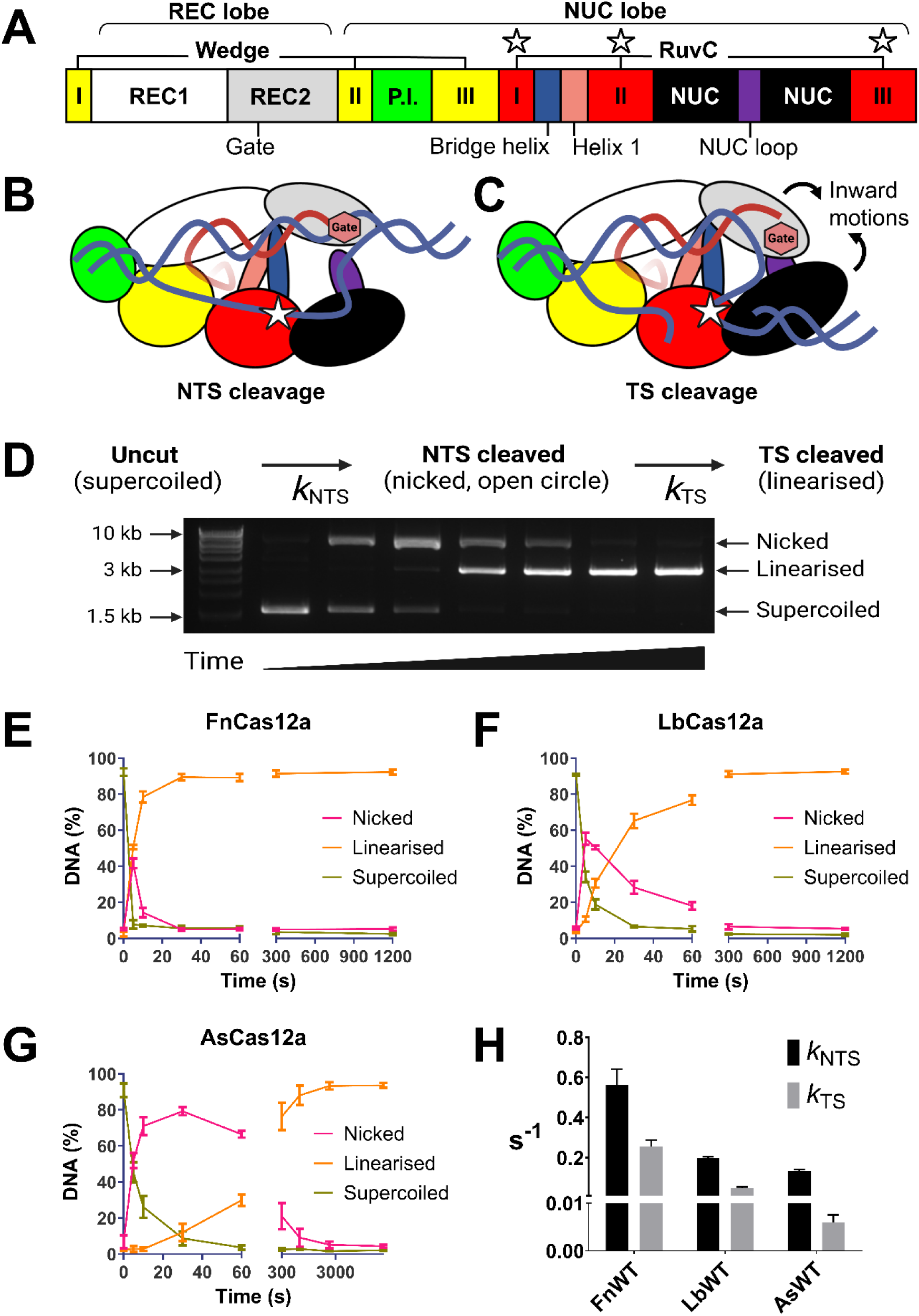
(**A**) General domain organisation of Cas12a orthologues (not to scale). Key residues and motifs are highlighted; the REC2 ‘gate’, the bridge helix and helix 1 -which comprise the ‘BH’ domain, the NUC loop (purple), and the RuvC active site residues (white stars). (**B,** C) Model of Cas12a tertiary structure bound to crRNA (red) and double-stranded DNA (blue). Depicted is the cis cleavage mechanism, where the NTS is cleaved first (**B**), followed by TS cleavage (**C**). Key residues and motifs in this mechanism are highlighted; REC2 ‘gate’ (red hexagon), NUC loop (purple), and RuvC active site (white star). (**D**) Schematic showing changes in plasmid DNA topology with sequential NTS and TS cleavage. Example agarose gel of plasmid cleavage over time (image over-exposed to show faint DNA bands). DNA cleavage by Cas12a results in evident changes in plasmid DNA topology, from the uncut and negatively supercoiled form (migrates at ∼1.5 kb), to the nicked open-circle form (migrates at ∼9 kb), to the linearised form (migrates at ∼3 kb). (**E, F, G**) Quantification of DNA fractions (nicked, linearised, supercoiled) over time, when incubated with (**E**) WT FnCas12a, (**F**) WT LbCas12a, and (**G**) WT AsCas12a – note longer time points. Line shows mean ± s.d. (**H**) Mean rate constants for NTS and TS cleavage (± s.d.) modelled from the change in DNA fractions over time, using data from (**E, F, G**).

The non-hybridised DNA strand (non-target strand, NTS) is coordinated close to the RuvC, and in the required 5’ to 3’ polarity for in-line nucleophilic attack by the RuvC; while the target strand (TS) is hybridised to the crRNA, stuck to the REC lobe far from the active site and in the opposite polarity (4, 13, 15, 16) (**Figure 1B, C**). Consequently, kinetic studies of sequential DNA strand cleavage have determined NTS cleavage is 2-20x faster than TS cleavage (14, 18, 20, 23, 26, 27). The NTS occludes the TS from the active site until it is cleaved, making an obligatory sequential cleavage mechanism of NTS cleavage preceding TS (13, 25) (**Figure 1B, C**).

Structures of Cas12a show the scissile phosphate of the TS is some 25 Å from the RuvC active site (13–16). Dynamic ‘pinching’ motions of the REC2 and NUC lobes have been observed in single-molecule FRET studies and molecular dynamics simulations (14, 24, 28–31), motions which shorten the distance required for the TS to traverse. A key aromatic ‘gate’ residue in the REC2 stacks at the 20^th^ position of the crRNA-TS heteroduplex, regulating the length of the R-loop and constraining the flexible fraying of the 3’ R-loop junction (25, 26) (**Figure 1B, C**). After *cis* cleavage, Cas12a remains stably bound to the PAM-proximal fragment of dsDNA containing the entire 20bp crRNA:TS heteroduplex used for target recognition (19). This ternary complex remains catalytically active to cleave nucleic acids in *trans* (9, 10).

This mechanism of *cis* and *trans* cleavage is considered to be consistent across Cas12a orthologues. Yet, when it comes to applying these enzymatic capabilities to genome editing and molecular detection, efficacies vary greatly between orthologues. These differences raise the question – what makes an effective Cas12a in which context? In a time when abundant Cas12a genes can be obtained from sampling environmental DNA, from genetic databases, and even designed *de novo*, resolving this question would greatly improve the efficiency of finding or engineering improved Cas12a nucleases.

To explore this question, we undertook a comparative study of three well-characterised Cas12a orthologues, from *Acidaminococcus sp. BV3L6* (AsCas12a), *Lachnospiraceae bacterium ND2006* (LbCas12a), and *Francisella tularensis subsp. novicida U112* (FnCas12a). When compared side-by-side in human cell lines, editing by AsCas12a and LbCas12a is more robust than FnCas12a, across a range of PAMs and target sites (5, 6). For *trans* cleavage activity, it is LbCas12a that has more robust activity than AsCas12a and FnCas12a (9, 10, 32).

AsCas12a, LbCas12a, and FnCas12a have high structural similarity (<3 Å RMSD) with less than 50% sequence similarity (4, 13–18). Given their shared mechanism of target DNA cleavage, they each ‘solve’ the same molecular problem with somewhat different amino acid sequences. We suspected this divergence causes their different performance in applied settings.

To interrogate this hypothesis, we generated mutations in the REC2 and NUC domains to explore what drives their DNA cleavage kinetics. We identified an uncharacterised ‘NUC loop’ as a structural element that traverses the distance between NUC and REC2 domains. This loop is present across Cas12a orthologues but varies considerably in length and amino acid sequence. Furthermore, the NUC loop is unresolved in most experimentally determined structures of Cas12a, suggesting it may be highly dynamic. Only recently have cryo-EM structures captured the NUC loop in the process of coordinating DNA strands for cleavage (18).

Across FnCas12a, LbCas12a, and AsCas12a, we thoroughly characterise REC2 and NUC loop mutants for their *cis* and *trans* cleavage kinetics, and their ability to interfere with plasmid transformation in *E. coli* and edit genes in human cell lines. We find apparent trade-offs between NTS, TS, and *trans* cleavage, which are driven by REC2 ‘gate’ and NUC loop interactions. Although mutagenesis could modulate *cis* cleavage rates 5-fold, there remained very large differences in between Cas12a orthologues, with FnCas12a displaying extremely rapid and robust DNA cleavage. To resolve this conundrum, and elucidate the dynamic role of the NUC loop, we conducted molecular dynamics simulations to compare the properties of REC2 – NUC dynamics between Cas12a orthologues. Together with recent cryo-EM and molecular dynamics (MD) results, this simulation shows the NUC loop makes dynamic interactions with the REC2 and the crRNA-TS heteroduplex. Furthermore, we found large differences in REC2 – NUC distance distributions, which may underwrite their different efficiencies of allosterically activating DNA catalysis. In total, this work advances our understanding of the mechanisms of nuclease activities of Cas12a orthologues.

## Materials and Methods

### Cloning

The coding sequences of WT FnCas12a and AsCas12a were cloned from parent vectors (Addgene #90094 and #90095 respectively) into a pET21_6His_2NLS vector that was a kind gift of Wolfe Lab (Addgene #114366), as described previously (21). Similarly, plasmid interference ‘locus’ plasmid was cloned with the In-Fusion kit (Takara Bio), as previously described (21).

Mutant sequences, such as Cas12a mutants and interference assay off-target plasmid, were generated using the Q5 site-directed mutagenesis kit according to the manufacturer’s instructions (New England Biolabs – NEB), and sequences verified with Sanger sequencing (Biomolecular Resource Facility, ANU) (**Table S5** – primers, **Table S6** – ‘locus’ oligonucleotides).

### Protein purification

All WT and mutant Cas12a proteins were purified with the following protocol. Plasmids were transformed into T7-express chemically competent cells (NEB), colonies picked, and small volumes (∼5 mL) grown overnight in Luria Broth supplemented with 100 µg/ml Ampicillin. Overnight cultures were used to inoculate a 250 mL culture, grown at 37 °C for ∼2 h in baffled flasks and vigorous shaking, until OD_600_ ∼0.6. Flasks were put on ice for 30-45 mins before addition of 1 mM IPTG, then transferred to an 18 °C incubator for shaking at 200 rpm overnight.

Expression cultures were centrifuged for 10 mins at 5,000 g, and pellets resuspended in Lysis Buffer (50 mM Tris-HCl pH 7.5, 500 mM NaCl, 5% glycerol, 1 mM DTT), with addition of one ‘cOmplete protease inhibitor tablet’ (Roche) per 50 mL of resuspension. Cells were lysed with sonication (Branson Sonifier), and supernatant clarified with 2 x 30 min centrifuge spins at 13,500 g. We performed metal affinity chromatography by loading the supernatant on an equilibrated Ni-NTA HisTrap column (GE Healthcare, 5 mL) with an AKTA Explorer (GE Healthcare), washed with Buffer A (50 mM Tris-HCl pH 7.5, 500 mM NaCl, 20 mM imidazole, 5% glycerol), and eluted with a stepwise addition of Buffer B (50 mM Tris-HCl pH 7.5, 500 mM NaCl, 500 mM imidazole, 5% glycerol). Fractions were analysed with SDS-PAGE, and peak fractions containing Cas12a were pooled, and diluted 2.5x with diluting buffer (50 mM Tris-HCl pH 7.5, 5% glycerol) to achieve 200 mM NaCl for cation exchange chromatography. After loading on a HiTrap Heparin column (GE Healthcare, 5 mL) pre-equilibrated with Buffer H-A (50 mM Tris-HCl pH 7.5, 200 mM NaCl, 5% glycerol), elution was performed with a linear gradient of Buffer H-B (50 mM Tris-HCl pH 7.5, 1 M NaCl, 5% glycerol). Again, fractions containing Cas12a were determined by SDS-PAGE and concentrated to a small volume (∼500 µL) by centrifugal molecular-weight cut-off tubes (30kDA, Pierce ThermoFisher). Concentrated protein was then buffer-exchanged into storage buffer (50 mM Tris-HCl pH 7.5, 500 mM NaCl, 50% glycerol, 1 mM DTT) with 0.5 mL centrifugal molecular-weight cutoff tubes (Millipore). Protein concentration was estimated with a Nanodrop spectrophotometer (ThermoFisher) using extinction coefficients calculated with Expasy ProtParam(33) and stored at -20 °C. Yields of Cas12a varied from 2-8 g/L of expression culture.

### Protein thermostability assay

Melt curves were conducted in 40X SYPRO Orange dye (ThermoFisher), and a StepOnePlus qPCR machine (Applied Biosystems), following the manufacturers detailed protocol (ThermoFisher). Cas12a proteins (in storage buffer) were diluted in nuclease-free water (Ambion) to a concentration of ∼1 µg per well, 3 x 20 µL replicates were added to a MicroAmp Fast 96-well Reaction Plate (Applied Biosystems), and fluorescence monitored at 1 °C increments from 25 to 99 °C. Melting temperature was defined as the fluorescence peak.

### Cis cleavage kinetics

In this assay, Cas12a was complexed with a crRNA targeting a site in a negatively supercoiled plasmid. RNAs were ordered from Integrated DNA Technologies (IDT) and resuspended in IDTE buffer (IDT) (**Table S7** – crRNA sequences). Cas12a-crRNA complexes were assembled by incubation at 25°C for 10 mins. Complexes were diluted in 1x cleavage buffer (10 mM Tris-HCl, pH 7.5, 10 mM MgCl2, 50 mM NaCl, 5 µg/ml BSA, 0.1 mM DTT) to a final concentration of 100 nM, and equilibrated at 30 °C on a thermocycler block prior to addition of target DNA.

‘DNA solution’ containing target plasmid DNA was diluted in 1x cleavage buffer to a final concentration of 10 nM, and also equilibrated at 30 °C. Equal volumes (5 µL) of Cas12a complex and DNA solution were mixed and incubated for the set time-points, and reaction quenched with addition of 5 µL ‘STOP solution’ (15%v/v proteinase K (Bioline), 250 mM EDTA, 50% v/v 5x loading dye (Bioline), in nuclease-free water (Ambion)). To remove DNA-bound Cas12a ribonucleoprotein, all samples were incubated at 55°C for 30 mins after addition of STOP solution.

DNA products were separated by gel electrophoresis (100V for 40 mins) on a 1.5% agarose gel pre-stained with 0.5x GelRed (Biotium). Gels were imaged with a Quantum geldoc (Vilber) with short exposure times to avoid oversaturation, and DNA band intensity quantified with ImageJ (34) (NIH). Changes in target plasmid topology were validated by nuclease digestion by Nt.BspQI and EcoRI (NEB), using the manufacturers protocol. Pixel numbers from DNA fractions ‘nicked’, ‘linear’, and ‘supercoiled’ were summed (Microsoft Excel), from which was calculated the percentage of nicked/linear/supercoiled DNA fractions. Percentage of DNA fraction at time points were used as input for kinetic modelling, where each replicate was individually modelled. Three replicates were performed for each Cas12a nuclease.

### Modelling rates of sequential DNA strand cleavage

The rate of change between DNA fractions was then modelled to obtain *k*_obs_ for both NTS and TS cleavage, where ka = *k*_NTS_ and kb = *k*_TS_. Rates were modelled in Berkeley Madonna (35), using the following equations, as previously detailed in the literature (26, 27, 36).

d/dt (SC) = -ka*SC - kini*SC

d/dt (ucSC) = kini*SC

d/dt (NICK) = ka*SC - kb*NICK - kini2*NICK -ka*NICK1

d/dt (NICK1) = -ka*NICK1 - kini*NICK1

d/dt (ucNICK) = kini2*NICK + kini*NICK1

d/dt (LIN) = kb*NICK

TotSC = SC + ucSC

TotNICK = NICK + NICK1 + ucNICK

Init SC = input initial %SC from dataset

init LIN = input initial %LIN from dataset

init NICK1 = input initial %NICK from dataset

init NICK = 0

init ucSC = 0

init ucNICK = 0

ka = 0.50

kb = 0.50

kini = 0.01

kini2 = 0.01

Where variables “SC”, “NICK”, and “LIN” were fitted to their corresponding dataset, being the percentage of nicked/linear/supercoiled DNA at timepoints.

### Trans cleavage assays

Cas12a RNP complexes were prepared as described for *cis* cleavage (10 nM final concentration), and *trans*-active complexes were made by addition of 1 nM target-strand DNA (**Table S6**) in 1 x cleavage buffer, followed by incubation at 30 °C for 45 mins. This active complex was then further diluted in 1x cleavage buffer, and 50 µL added to wells of a flat-clear-bottom black fluorescence 96-well plate (ThermoFisher). Fluorescent-quencher reporter ssDNA (**Table S6**) was prepared in 1x cleavage buffer to a final concentration of 75 nM, 50 µL of which was added into each well, using the dispenser pump of a Victor Nivo plate reader (PerkinElmer). Excitation and emission filters of 480/30nm and 530/30nm respectively were used to measure fluorescence over time. Three replicates were performed per condition.

### Structural analysis

Protein Data Bank (PDB) files for the accession codes indicated were obtained from PDB.com, and structural alignment performed in VMD (37), using the STAMP algorithm (38). Predicted Cas12a structures were obtained from Alphafold/EMBL (39, 40), to fill structural elements not modelled in experimental structures of AsCas12a and LbCas12a. UniProt identifiers were U2UMQ6 for AsCas12a, and A0A182DWE3 for LbCas12a.

### Plasmid interference assay

This assay used three plasmids, one expressing Cas12a under a T7 promoter (AmpR), another encoding a crRNA under a T7 promotor (CmR) with a spacer sequence matching a third ‘target’ plasmid (KanR).

To achieve high rates of plasmid transformation, chemically competent T7Express cells (NEB) were made harbouring both the Target plasmid, and either the crRNA expressing (+ crRNA) or empty vector (no crRNA). 20 ng of Cas12a plasmid was transformed into these +/- crRNA strains by heat shock at 42°C and recovered with SOC media at 37°C for 30 mins. Serial dilutions were plated onto triple selective media (100 µg/mL ampicillin, 50 µg/mL kanamycin, 25 µg/mL chloramphenicol) containing 0.5 mM IPTG, Plates were incubated overnight at 37°C for approximately 16 h, and colonies counted. Three replicates were performed for each transformation. Statistical significance was calculated by two-way ANOVA followed by Tukey’s multiple comparisons test, using GraphPad Prism 10 (GraphPad Software, www.graphpad.com).

### Human cell line editing

Human genome targets with previously identified off-target sites were chosen for gene editing experiments (6, 7, 41). The target sites respectively have the canonical PAM motifs, TTTA, TTTC, and TTTG, to minimise PAM bias in editing efficiencies between Cas12a orthologues (6, 16, 42–44). The most-represented off-target sites identified by GUIDE-seq and DIGENOME-seq were chosen for high-throughput sequencing (6, 7, 41). crRNAs were ordered as HPLC-purified RNA (IDT) (**Table S7**).

HEK293T, A549, and Jurkat cell lines were obtained from the American Type Culture Collection (ATCC) and tested free of mycoplasma infection. HEK293T were cultured in high glucose Dulbecco’s Modified Eagle Medium (Gibco) supplemented with 10% FBS and 1x Penicillin-Streptomycin-Glutamine (Gibco). Jurkat cells were cultured in RPMI-1640 supplemented with 10% FBS and 1x Penicillin-Streptomycin-Glutamine (Gibco). A549 were cultured in Ham’s F-12K (Kaighn’s) Medium (Gibco) supplemented with 10% FBS and 1x Penicillin-Streptomycin (Gibco). Cells were maintained at 37 °C, with 5% CO2 in a humidified atmosphere, and transfected at passage 10.

Cas12a proteins were assembled with their cognate crRNA, targeting either DNMT1-3, DNMT1-7, or AGBL1. Each RNP reaction consisted of 0.575 µM crRNA with 32 pM of Cas12a. The crRNA and Cas12a were complexed in 2.2 µL Neon Transfection System ‘R’ resuspension buffer (Invitrogen) at 37 °C for 5 mins and left at room temperature post-complexing.

Electroporation was conducted using Neon Transfection System (Invitrogen) according to the manufacturer’s protocol, with the following modification, that all three cell lines were resuspended in Neon Transfection System ‘R’ resuspension buffer (Invitrogen) to a concentration of 1×10^7^/ml.

For each electroporation reaction, 1×10^5^ cells prepared above, were incubated with 1xRNP at 37 °C for 5 mins, before being electroporated using the 10 µL Invitrogen Neon Transfection System. Electroporation protocols for cell lines were HEK293T; 1,150 volts, 20 ms, 2 pulses; Jurkat; 1,325 volts, 10 ms, 3 pulses; A549; 1,230 volts, 30 ms, 2 pulses. Two reactions were seeded per well, in a 24-well plate. Cells were recovered in complete medium at 37 °C with 5% CO2 for 72 h. Controls for each cell line included no electroporation, and electroporation sans nuclease. Three replicate were performed for each control and Cas12a.

Samples were harvested at 72 hours post transfection, including growth media to capture all cells, dead or alive. Cells were pelleted at 500 g for 2 mins at room temperature, then washed with 1x PBS. Cells were again centrifuged at 500 g for 2 mins at room temperature, PBS was removed, and samples were then stored at -20 °C prior to DNA extraction. Samples were thawed on ice and genomic DNA was extracted using the ISOLATE II Genomic DNA Kit (Meridian Bioscience), and following the manufacturer’s instructions, with the sole modification of eluting twice in with the same 100 µL of elution buffer.

### Quantification of genome editing by high-throughput sequencing

Primers were designed for high-throughput sequencing of identified on and off-target sites of DNMT1-3, DNMT1-7, and AGBL1; positioned ∼125 bp upstream and downstream of the target site, resulting in ∼250 bp amplicon (primers in **Table S8**). Primers and target DNA were dispensed in 384 well plates, and Illumina Ampliseq used to perform a paired-end 250 bp library preparation.

Quality control was performed on the ∼32 million reads obtained, using FastQC (45). The reads were then analysed by CRISPResso2(46), using the Hg38 human genome as reference using the following parameters: --cleavage_offset 1 --quantification_window_size 20 -- ignore_substitutions --default_min_aln_score 50. Samples with less than 1,000 mapped reads were discarded. Insertions and deletions in this window were combined to calculated total percentage of indels at a given on or off-target site. Statistical significance was calculated by two-way ANOVA followed by Tukey’s multiple comparisons test, using GraphPad Prism 10 (GraphPad Software).

### Structural models for simulation

Molecular simulations were based on three structures of Cas12a across different species as obtained post *cis*-cleavage: (1) the cryo-EM structure of FnCas12a (PDB: 6GTG) (14) at 2.50 Å resolution, (2) the X-ray structure of AsCas12a from (PDB: 5B43) (15) at 2.80 Å, (3) the X-ray structure of LbCas12a (PDB: 5XUS (16) at 2.5 Å. All systems were embedded in explicit waters and counterions were added to neutralize the total charge, leading to periodic cells of ∼138×149×167 Å^3^ and ∼307,000 atoms for each system.

### Molecular dynamics simulations

Molecular Dynamics (MD) simulations were performed using a protocol tailored for RNA/DNA nucleases using the Amber ff19SB force field (47), including the ff99bsc1 corrections for DNA (48), and ff99bsc0+χOL3 corrections for RNA (49, 50). The TIP3P model was employed for explicit water molecules (51), and the Li & Merz 12-6 model of non-bonded interactions was used for Mg^2+^ ions (52). We have extensively employed these force field models in computational studies of CRISPR-Cas systems (29), showing also that they perform well for long timescale simulations (24). The Li & Merz model also reported a good description of Mg^2+^ bound sites, in agreement with quantum/classical simulations (53). An integration time step of 2 fs was employed. All bond lengths involving hydrogen atoms were constrained using the SHAKE algorithm (54). Temperature control (300 K) was performed via Langevin dynamics (55), with a collision frequency γ = 1. Pressure control was accomplished by coupling the system to a Berendsen barostat (56), at a reference pressure of 1 atm and with a relaxation time of 2 ps.

The systems were subjected to energy minimization to relax water molecules and counterions, keeping the protein, the RNA, DNA and Mg^2+^ ions fixed with harmonic position restraints of 300 kcal/mol · Å^2^. Then, the systems were heated up from 0 to 100 K in a canonical ensemble (NVT), 120 by running two simulations of 5 ps each, imposing position restraints of 100 kcal/mol · Å^2^ on the above-mentioned elements of the system. The temperature was further increased up to 200 K in ∼100 ps of MD in the isothermal-isobaric ensemble (NPT), reducing the restraint to 25 kcal/mol · Å^2^. Subsequently, all restraints were released, and the temperature of the systems was raised up to 300 K in a single NPT simulation of 500 ps. After ∼ 1.1 ns of equilibration, ∼10 ns of NPT runs were carried out allowing the density of the systems to stabilize around 1.01 g cm^-3^. Finally, production runs were carried out in the NVT ensemble in 4 replicates, collecting ∼1 μs for each replicate. These simulations were performed using the GPU-empowered version of AMBER 20.

For distance and contact analyses, domains were defined as follows: FnCas12a, REC2 (340-591) and NUC (1079-1254), LbCas12a, REC2 (283-521) and NUC (998-1179), and AsCas12a REC2 (321-526) and NUC (1067-1262). Distance analysis considered the centre of mass of the selected regions, and a contact was considered when the distance between two heavy atoms among the regions of interest was less than 3.5 Å. Amino acids involved in contacts were identified by visual inspection of trajectories. Kernel Density Estimation (KDE) plots were employed to visualize the probability density of REC2-NUC distances and contacts, using the ‘kdeplot’ function from the seaborn library, a statistical data visualization package in Python (57).

## Results

### Cas12a orthologues display distinctly different rates of sequential strand cleavage

Central to the natural or applied functions of Cas12a is *cis* cleavage. The kinetics of sequential DNA strand cleavage have been determined for wild-type FnCas12a, LbCas12a, and AsCas12a, and have yielded rate constants that vary by several orders of magnitude (**Table S1** (14, 18, 20, 23, 26–28)). This can be attributed to a number of factors known to affect *cis* cleavage kinetics; temperature (58), DNA substrate topology (27), and magnesium ion concentration (20, 28). Thus, we wished to undertake kinetic comparisons in conditions at which each three orthologues had previously been determined to be maximally active (59); 30°C, pH 7.5, 50 mM NaCl (plus 10 mM Tris-HCl, 10 mM MgCl_2_, 5 µg/mL BSA, 0.1 mM DTT).

We determined the *cis* cleavage kinetics of the three Cas12a orthologues using a plasmid cleavage assay developed by the Szczelkun Lab (26, 27, 36). Briefly, the sequential DNA strand cleavage of a negatively supercoiled plasmid causes sequential changes in plasmid DNA topology, transitions that are visible in gel electrophoresis (60) (**Figure 1B, C, D, Figure S1**). NTS cleavage relaxes the supercoiled plasmid into the open-circle form, and TS cleavage converts the open-circle to the linearised form (26, 27, 36). Quantification of these topological changes over time allows modelling of strand cleavage rates (26, 27, 36).

In this time-course assay of plasmid DNA cleavage, FnCas12a exhibited the fastest DNA cleavage, with the fraction of linearised DNA plateauing at 30 seconds (**Figure 1E**). LbCas12a linearised the target plasmid by 300 seconds (**Figure 1F**), and AsCas12a by the 2,700 second time-point (45 minutes, **Figure 1G**). Fitting a sequential-strand cleavage model to this time-course assay data yielded observed rate constants for NTS and TS cleavage (26, 27, 36).

This modelling showed the large time differences in linearising plasmid DNA are driven by very different *cis* cleavage kinetics (**Figure 1G**). FnCas12a exhibited a *k*_NTS_ 2.8x faster than LbCas12a, which in turn had a *k*_NTS_ 1.4x faster than AsCas12a (**Figure 1G**, **Table S1**). The differences were greater with *k*_TS_; FnCas12a was 5.2x faster than LbCas12a, which was 8.2x faster than AsCas12a (**Figure 1G**, **Table S1**).

This order of *cis* cleavage speed, where FnCas12a > LbCas12a > AsCas12a, is evident in other reports (comparing FnCas12a, LbCas12a, and AsCas12a (19); and between FnCas12a and AsCas12a (25)). Of these three orthologues, FnCas12a is generally reported to have lower gene editing efficiency in human cell lines, and weaker *trans* cleavage (5, 6, 9, 10, 32). We therefore expected FnCas12a to have a defect in *cis* cleavage. But in the conditions tested, FnCas12a has the most robust *cis* cleavage. This raises the question; what drives the different *cis* cleavage kinetics of Cas12a orthologues, and how does it relate to genome editing and DNA detection?

### REC2 domain mutations reduce NTS and trans cleavage rates

The NTS cleavage mechanism of Cas12a is straightforward; a groove of DNA binding residues across the Wedge, RuvC, and NUC domains guide the NTS into the RuvC active site in the correct 5’ to 3’ polarity for in-line nucleophilic attack (4, 13, 14, 18). The mechanism of TS cleavage is less simple, the scissile phosphate must traverse over 20 Å and twist 180° to enter the RuvC with the correct polarity (4, 13, 25, 26). This conformation of the TS is allowed by unwinding at the 3’ end of the crRNA:TS R-loop (14, 18, 25, 26).

A ‘gate’ residue in the REC2 plays a key role in this process, stacking after the 20^th^ position of the crRNA:TS heteroduplex and regulating the length of the R-loop (4, 13–16, 18, 26). Removing this stacking interaction by alanine substitution increased TS cleavage rates in LbCas12a (26). Despite increased *k*_TS_, this mutant displayed slower *trans* cleavage (61, 62).

We were intrigued by this apparent trade-off between TS cleavage and *trans* cleavage. To explore this, we generated alanine substitutions of the REC2 gate for three Cas12a orthologues and assayed their *cis* and *trans* cleavage kinetics.

The time-course plasmid cleavage assay was performed for FnY410A, LbW355A, and AsW382A (**Figure S2**). Consistent with previous reports, *k*_TS_ increased for all three mutants (**Figure 2A, B, C**) (26). Interestingly, this came at a cost of decreased *k*_NTS_. This effect was remarkably similar across all three alanine substitution mutants, where *k*_NTS_ decreased by 1.5 to 2.5x, and *k*_TS_ increased by 3.3 to 4.7x (**Table S2**). This suggests the two strands are in competition for access to the RuvC active site.

**Figure 2:**
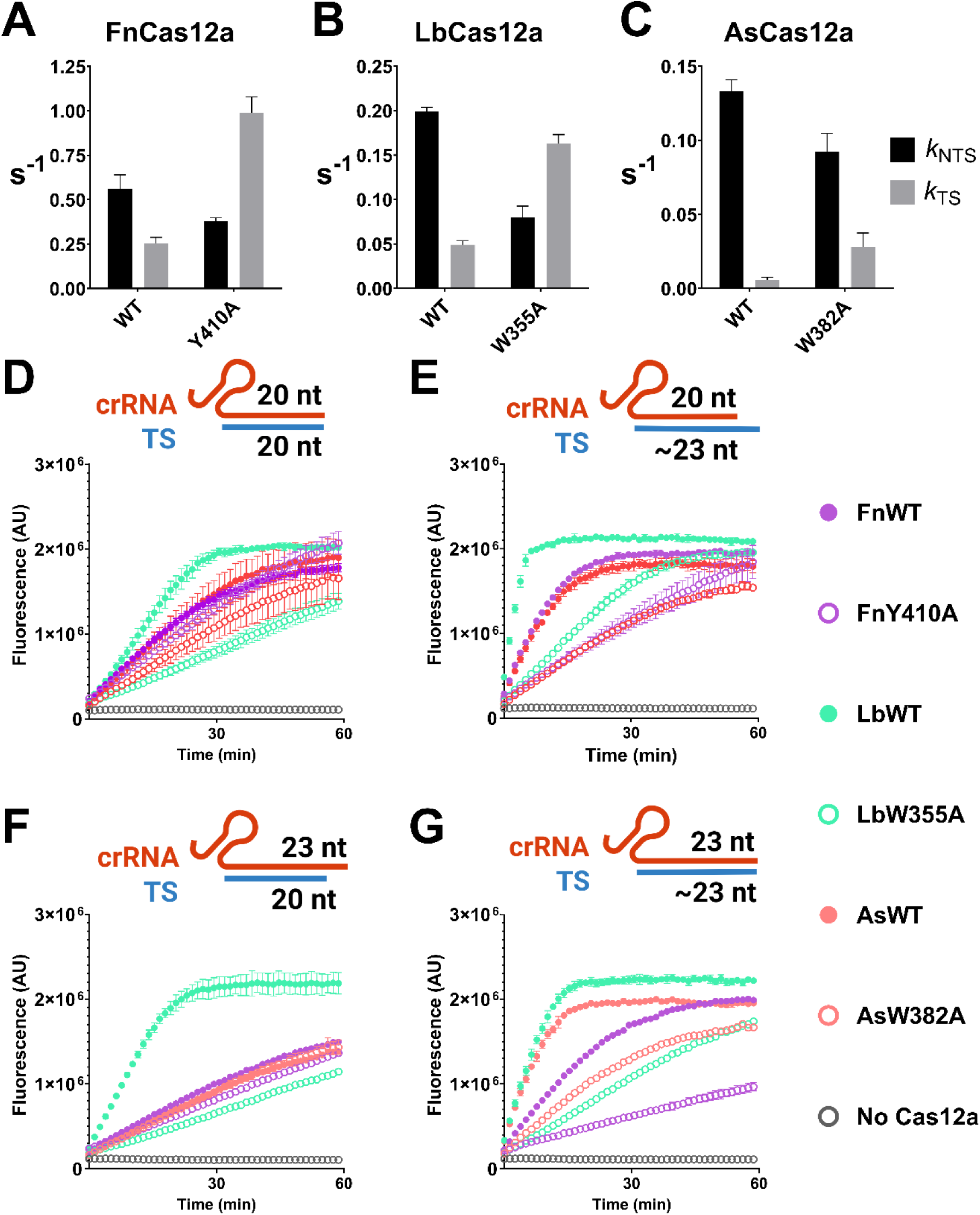
Mean rate constants for NTS and TS cleavage (± s.d.) for REC2 mutants of (**A**) FnCas12a, (**B**) LbCas12a, (**C**) and AsCas12a. (**D, E, F, G**) Trans cleavage curves of WT and REC2 mutants, when activated with the combination of crRNA and TS indicated. Dots show mean ± s.d.

Having replicated the faster *k*_TS_ of REC2 ‘gate’ mutants, we aimed to test their *trans* cleavage kinetics. If the motion of ‘TS-loading’ can sterically hinder *trans* ssDNA substrates, then REC2 mutants should have slower *trans* cleavage than WT, but not in the case of a truncated target-strand.

To test this hypothesis, we assembled Cas12a-crRNA complexes with target strand ssDNA that was either full-length, or truncated at the 20^th^ position. These ‘TS-loading/*trans*-active’ complexes were made by a 45-minute *cis* cleavage reaction. In this experiment, the full-length target strand would be cleaved and trimmed to ∼22 - 24 nucleotides, depending on the orthologue (20, 22, 22, 25, 26).

We first compared Cas12a ternary complexes with a 20nt crRNA spacer sequence and truncated TS (**Figure 2D**). In this condition, FnY410A and AsW382A had similar rates of *trans* cleavage activity to their WT enzyme, while LbW355A had much slower rates than LbWT (**Figure 2D**). However, when complexed with a full-length TS, REC2 mutants had consistently slower *trans* cleavage activity compared to WT (**Figure 2E**).

Previous work has also demonstrated that 3’ extension of the crRNA past the 20^th^ position also influences *cis* and *trans* cleavage rates (10, 26, 63). We repeated the truncated/full-length target strand comparison, but with crRNAs consisting of a 23nt spacer sequence. We observed a similar pattern as with 20nt crRNAs; where REC2 mutants have similar-to-WT rates of *trans* cleavage with the truncated TS (LbW355A again the exception, **Figure 2F**), and much decreased *trans* cleavage with the full-length TS (**Figure 2G**).

Overall, these data suggest that TS-loading can indeed slow *trans* cleavage. This agrees more broadly with data showing competition at the RuvC active site; namely that NTS-loading slows TS cleavage, and that excess ssDNA can slow TS cleavage (25, 28). It appears likely that NTS, TS, and *trans* DNA strands compete to be coordinated in the narrow RuvC active site.

Previous work showed the AsW382A mutant had decreased gene editing activity in human cell lines (15), and lesser stability of LbW355A was inferred from breakdown products seen in SDS-PAGE (26). We tested WT and REC2 mutants for their ability to interfere with plasmids in *E. coli*, and found no defect in their function (**Figure S3A, B**). Furthermore, we tested protein thermostability (64), and found no defect in the stability of WT or REC2 mutant *apo* proteins (**Figure S3C**).

### NUC loops are critical to Cas12a function

The REC2 ‘gate’ appears to regulate *k*_TS_, yet there remains a ∼20 Å distance for the TS to traverse from REC2 to RuvC. Structural data at the time showed a lack of electron density between the REC2 and NUC lobes (4, 13–16) – with the exception of a single structure (14). A ‘transient state’ ternary structure of FnCas12a captured a loop extending from the bulk of the NUC lobe towards the 3’ end of the crRNA-TS heteroduplex (14). We hypothesised this NUC loop could be key in the TS-loading mechanism of Cas12a orthologues.

Sequence alignment showed the NUC loop has divergent amino acid composition across the three Cas12a orthologues (**Figure 3A**), and structural prediction shows different conformations (**Figure S4**). These loops contain a variety of charged and aromatic amino acid sidechains that could interact with nucleic acids. To disrupt these potential interactions, two general mutations were designed. Firstly, to remove any specific interactions by the NUC loop, but retain the steric bulk, the ‘head’ of the loop was substituted for a flexible linker motif of repeating glycine-serine residues. This was termed the ‘FLX’ substitution (**Table S3**). Secondly, to remove both steric bulk and any protein-nucleic acid interactions, the NUC loop was deleted - the ‘ΔLoop’ mutation (**Table S3**).

**Figure 3:**
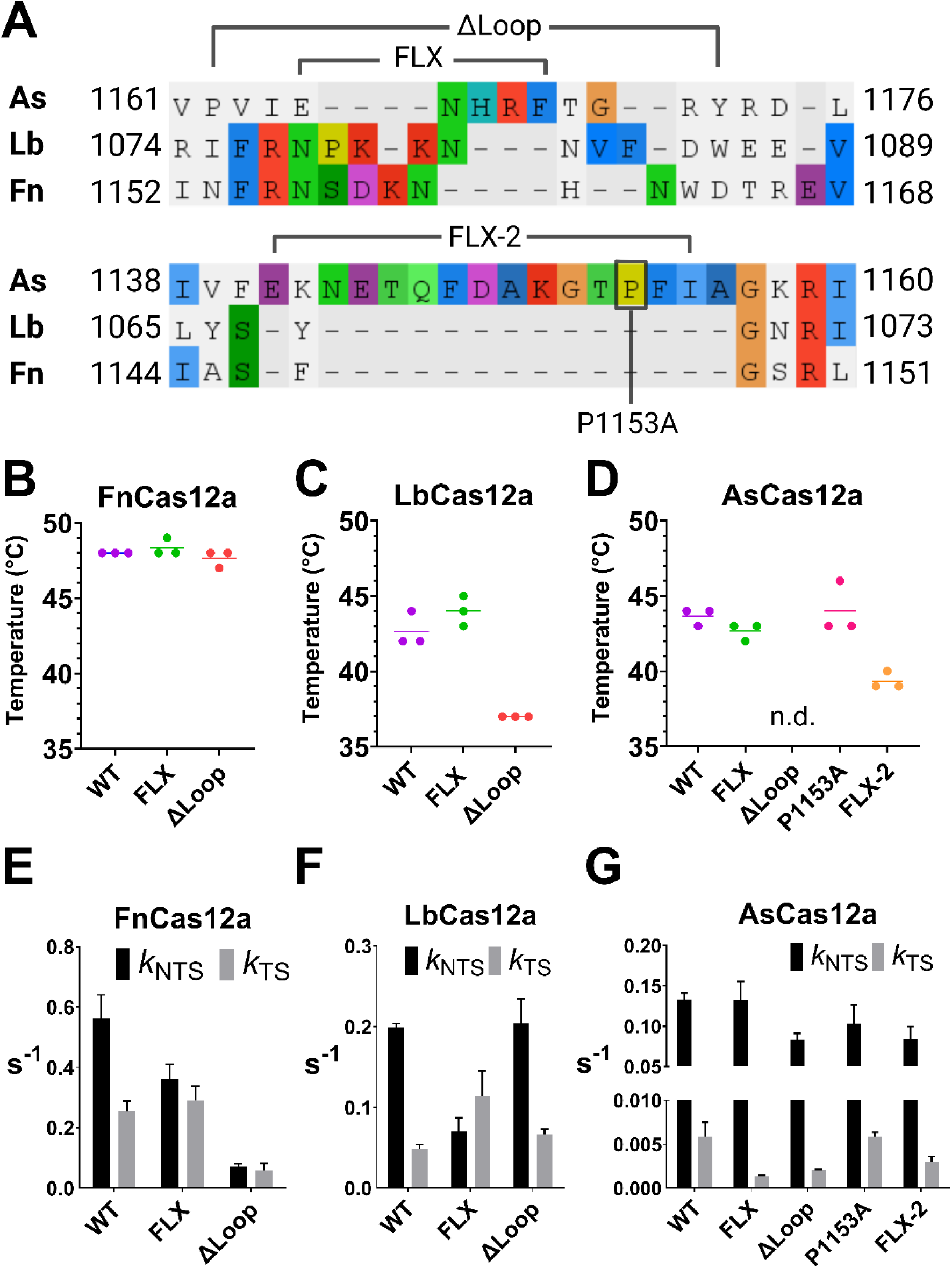
(**A**) Sequence alignment of selected region of the NUC domain of AsCas12a, LbCas12a, and FnCas12a, to illustrate the NUC loops. Mutations to disrupt the NUC loop are annotated; where ‘FLX’ indicates regions substituted for glycine-serine repeats, and ‘ΔLoop’ indicates regions deleted (**Table S3**). (**B, C, D**) Thermostability assay, showing melting temperature of three replicates, line shows mean. (**E, F, G**) Mean rates of NTS and TS cleavage (± s.d.), for WT Cas12a and NUC loop mutants as indicated.

Notably, AsCas12a has an additional insertion in the NUC lobe, not present in FnCas12a and LbCas12a, a motif we named ‘NUC loop 2’ (**Figure 3A, Figure S4**). This loop does not extend towards the heteroduplex, instead, it sits on the surface of the NUC and folds back towards the RuvC (**Figure S3**). Prolines often being crucial structural elements, we designed an alanine substitution of P1153 in NUC loop 2. Deletion of NUC loop 2 resulted in no soluble protein expression (data not shown), instead this motif was truncated and substituted with glycine-serine-glycine, a mutant we termed ‘FLX-2’ (**Table S3**).

We first tested the thermostability of the NUC loop mutants. FnFLX and FnΔLoop had thermostability similar to WT enzyme, each at ∼ 48 °C (**Figure 4A**). LbFLX had similar thermostability to LbWT, at 42 – 43 °C, while LbΔLoop was decreased at 37 °C (**Figure 4B**). AsFLX and AsP1153A were as stable as AsWT, at 43 – 44 °C, while AsFLX-2 was decreased at 39 °C, and AsΔLoop had no detectable fluorsecence peak (**Figure 4C**).

**Figure 4:**
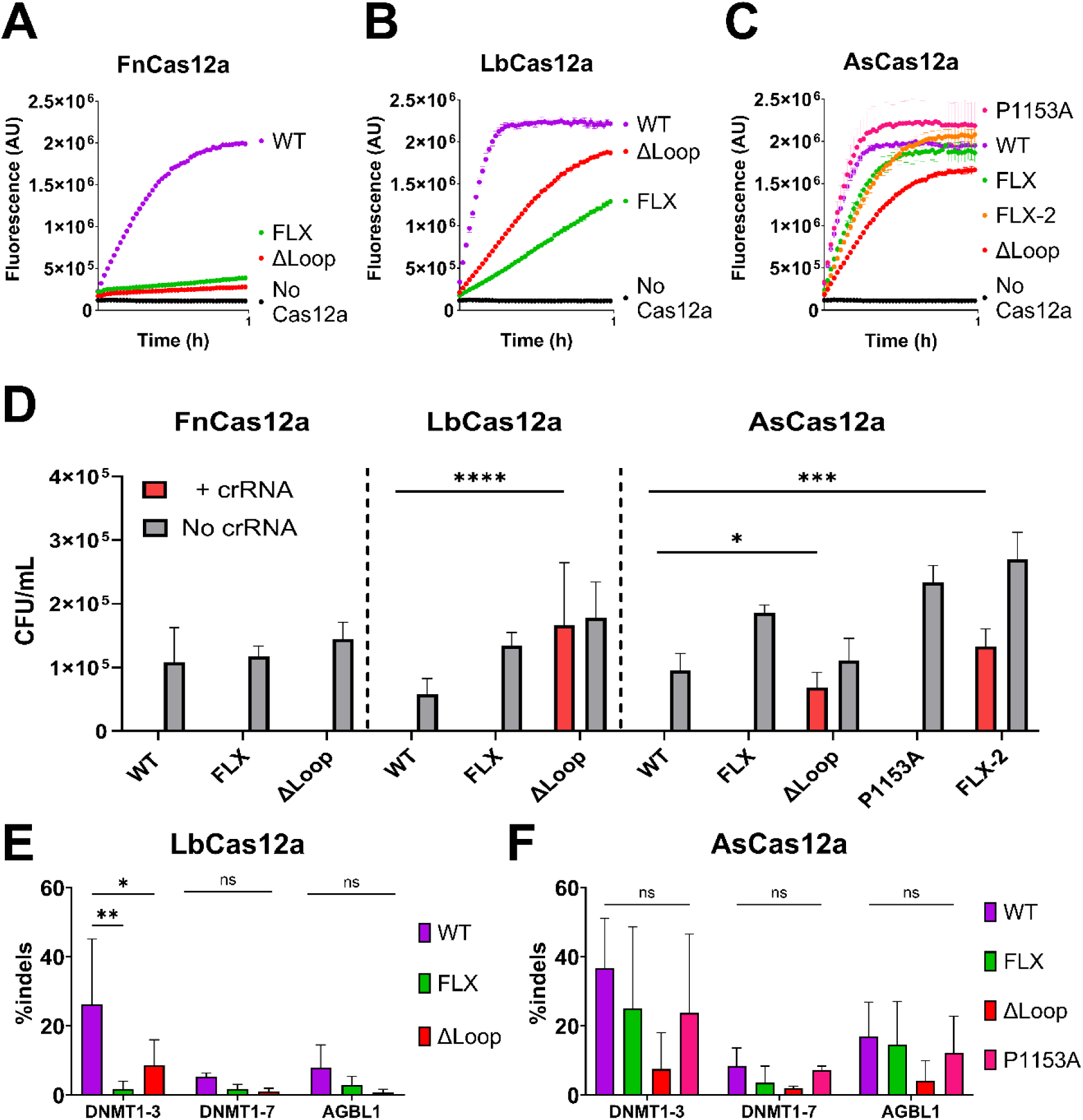
Trans cleavage curves of WT and NUC loop mutants of (**A**) FnCas12a, (**B**) LbCas12a, and (**C**) AsCas12a. Dots show mean, ± s.d. (**D**) Mean colony forming units per mL (error bars show s.d.), for ± crRNA conditions and NUC loop mutations as indicated. Statistical significance evaluated by two-way ANOVA with Tukey’s multiple comparison test (*p<0.1, **p<0.01, ***p<0.001, ****p<0.0001). Editing efficiency in HEK293T cell line. Mean percentage of insertions and deletions (indels) at the target site indicated (error bars show s.d.), for WT and NUC loop mutants of (**E**) LbCas12a, and (**F**) AsCas12a. Statistical significance evaluated by two-way ANOVA with Tukey’s multiple comparison test (*p<0.1, **p<0.01).

Next, we characterised the *cis* cleavage kinetics of NUC loop mutants (**Figure S5-78, Table S2**). The FnFLX substitution mutant had slower *k*_NTS_ by 1.5x, while *k*_TS_ increased by 1.1x (**Figure 3E**). A similar, but more pronounced effect was observed in LbFLX, with *k*_NTS_ decreased by 2.8x, while *k*_TS_ increased by 2.3x (**Figure 3F**). This is similar to alanine substitution of the REC2 ‘gate’, suggesting the NUC loop may regulate the order of strand cleavage for these orthologues. In contrast, AsFLX exhibited a very specific effect on *k*_TS_, decreasing rates by 6x, and leaving *k*_NTS_ essentially unchanged (**Figure 3G**).

NUC loop deletion had even more variable effects on DNA cleavage rates. FnΔLoop exhibited globally decreased *cis* cleavage compared to FnWT, *k*_NTS_ decreasing by 7.8x and *k*_TS_ decreasing by 4.2x (**Figure 3E**). Interestingly, FnΔLoop only linearised ∼75% of the plasmid target, leaving a large ‘nicked’ fraction (**Figure S5B**). This may indicate unstable target DNA binding. Despite its lesser thermostability, LbΔLoop showed similar *k*_NTS_ to LbWT, with *k*_TS_ slightly increased by 1.3x (**Figure 3F**). AsΔLoop decreased *k*_NTS_ by 1.6x, and halved *k*_TS_ relative to AsWT (**Figure 3G**).

Disruption of NUC loop 2 in AsCas12a decreased *cis* cleavage rates, despite its distance from any nucleic acids. P1153A substitution modestly decreased *k*_NTS_ by 1.3x, while *k*_TS_ was unchanged (**Figure 3G**). AsCas12a FLX-2 displayed very similar kinetics to AsΔLoop, *k*_NTS_ decreasing by 1.6x, and *k*_TS_ 3x slower (**Figure 3B**). These latter two mutants being the least thermostable of the mutants generated, their decreased *cis* cleavage may stem from globally disrupted protein function, rather than from loss of specific NUC loop interactions.

Overall, these data suggest integrity of the NUC loop is important for the *cis* cleavage activity of Cas12a orthologues. To further characterise the role of NUC loops, we assayed their *trans* cleavage activity, their plasmid interference in *E. coli*, and their editing activities in mammalian cell lines.

As with REC2 ‘gate’ mutants, the *trans* cleavage activity was tested with four different combinations of crRNA and target strand DNA. Unlike the REC2 mutants, NUC loop mutants did not show consistent patterns of substrate-dependant activity (**Figure S9-11**). For clarity, only *trans* cleavage reactions with the 23nt crRNA and full-length target strand are shown in **Figure 4**.

Despite their relatively robust *cis* cleavage, the FnFLX and FnΔLoop mutants had greatly decreased *trans* cleavage (**Figure 4A**). To a lesser extent, LbFLX and LbΔLoop mutants had reduced *trans* cleavage relative to LbWT (**Figure 4B**). AsP1153A showed similar *trans* cleavage to AsWT, while AsFLX, AsFLX-2, and AsΔLoop had moderately decreased *trans* cleavage activity (**Figure 4C**).

These data indicate the NUC loop plays a key role in the *trans* cleavage activity of FnCas12a. For LbCas12a and AsCas12a, NUC loop mutation decreases *trans* cleavage, although not to the magnitude of FnCas12a mutants.

Next, we tested the function of these NUC loop mutants for their ability to interfere with plasmid transformation in *E. coli*. Briefly, plasmids encoding Cas12a were transformed into *E. coli* strains harbouring either an empty vector (no crRNA) or a crRNA-encoding vector (+ crRNA) that directed Cas12a to cleave a third ‘target’ vector. FnFLX and FnΔLoop showed as robust plasmid interference as FnWT, with no colonies observed for the ‘+ crRNA’ condition (**Figure 4D**). LbWT and LbFLX also showed strong plasmid interference, but LbΔLoop showed a loss of activity, with CFU/mL counts similar to the ‘no crRNA’ transformations (**Figure 4D**). AsWT, AsFLX, and AsP1153A had robust plasmid interference, while AsΔLoop and AsFLX-2 lost plasmid interference activity (**Figure 4D**). Notably, it is the least thermostable Cas12a mutants (LbΔLoop, AsΔLoop, and AsFLX-2) that displayed the weakest plasmid interference, highlighting the importance of protein integrity in this assay.

As these NUC loop mutants had novel effects on target DNA cleavage *in vitro*, we aimed to assess their gene editing efficiency. Cas12a ribonucleoprotein complexes were electroporated into human cell lines, and insertions and deletions (indels) at target sites were quantified by high-throughput sequencing. Cas12a mutants were tested in HEK293T, A549, and Jurkat cell lines (**Figure 4E, Figure S13-15**), for clarity, only HEK293T editing is displayed in **Figure 4E**. We observed lower editing efficiency for FnCas12a compared to AsCas12a and LbCas12a (**Figure S12**), in agreement with previous works (5, 6). We therefore decided not to pursue further human cell line editing with FnCas12a mutants.

Despite the robust *E. coli* plasmid interference of LbFLX, it was significantly less active than LbWT in genome editing (**Figure 4E**). Expected from its lower activity in *E. coli*, LbΔLoop exhibited much lower editing compared to LbWT (**Figure 4E, Figure S13-15**). AsFLX and AsP1153A displayed a similar indel rate to AsWT, across cell lines and target sites (**Figure 4E, Figure S13-15**). AsΔLoop showed consistently decreased editing efficiencies, as expected from its weak interference in *E. coli*.

Given their slower cleavage of target DNA *in vitro*, we aimed to test if AsFLX and AsP1153A exhibited less off-target editing than AsWT. The most frequent off-targets for the DNMT1-3, DNMT1-7, and AGBL1 on-targets were derived from (6), and indels quantified by high-throughput sequencing (**Figures S16-18**). However, off-target editing with these nucleases was consistently low, with little difference between these nucleases and control electroporations (**Figures S16-18**).

Overall, the NUC loop can play a critical role in the *in vitro* and *in vivo* function of Cas12a, but this varies considerably between orthologues. Strikingly, although FnΔLoop was the most disrupted of all the FnCas12a mutants generated, it still retained *in vitro* TS cleavage rates 10x faster than wild-type AsCas12a. To understand this difference, we looked for more global drivers of catalytic function.

### Molecular dynamics simulations reveal distinct probabilities of ‘clamping’ between Cas12a orthologues

Recent works have demonstrated the importance of dynamic conformational changes in catalysis by Cas12a (14, 18, 24, 28–31, 65). On binding a matching DNA target, stable contacts are formed between the crRNA:TS heteroduplex and the REC2 domain (14, 18, 66). These interactions constrain the flexibility of the REC2 (18). This allows the BH domain to ‘dock’ with the REC2 domain and form contacts which ‘open’ the RuvC-lid, thus allosterically activating DNA cleavage (18, 21–23). In the process of sequentially cleaving NTS and TS, inward motions of REC2 – NUC domains have been observed in single-molecule FRET experiments, and predicted in molecular dynamics simulations (14, 24, 28–31).

The inward ‘clamping’ motions between REC2 and NUC are correlated with TS cleavage, in which an especially high FRET state is seen immediately before TS cleavage and substrate release (28, 30, 31). Given the disparities in *k*_TS_ we observe, we wished to compare REC2 – NUC dynamics between wild-type Cas12a orthologues. Furthermore, as the NUC loop extends towards the REC2 domain, we reasoned it may make contacts with the REC2 in the dynamic motions of DNA cleavage. To test this, we performed μs scale classical MD simulations of Cas12a orthologues. We employed structures of Cas12a in their post-*cis* cleavage state, which represents a TS-loading/ *trans*-active state.

This simulation detailed the residues involved in the REC2-NUC contacts (**Figure 5A**, enumerated in **Supp table S4**). Notable amongst these contacts is the NUC-loop. Additionally, the PAM-distal tip of the NUC lobe also makes contacts with the REC2. These NUC regions interact with the REC2 domain, in the most distal regions where the ‘gate’ residue is located. Notably, extensive NUC-loop and heteroduplex interactions were also seen in recent high-resolution simulations of FnCas12a cleaving the TS (24).

**Figure 5:**
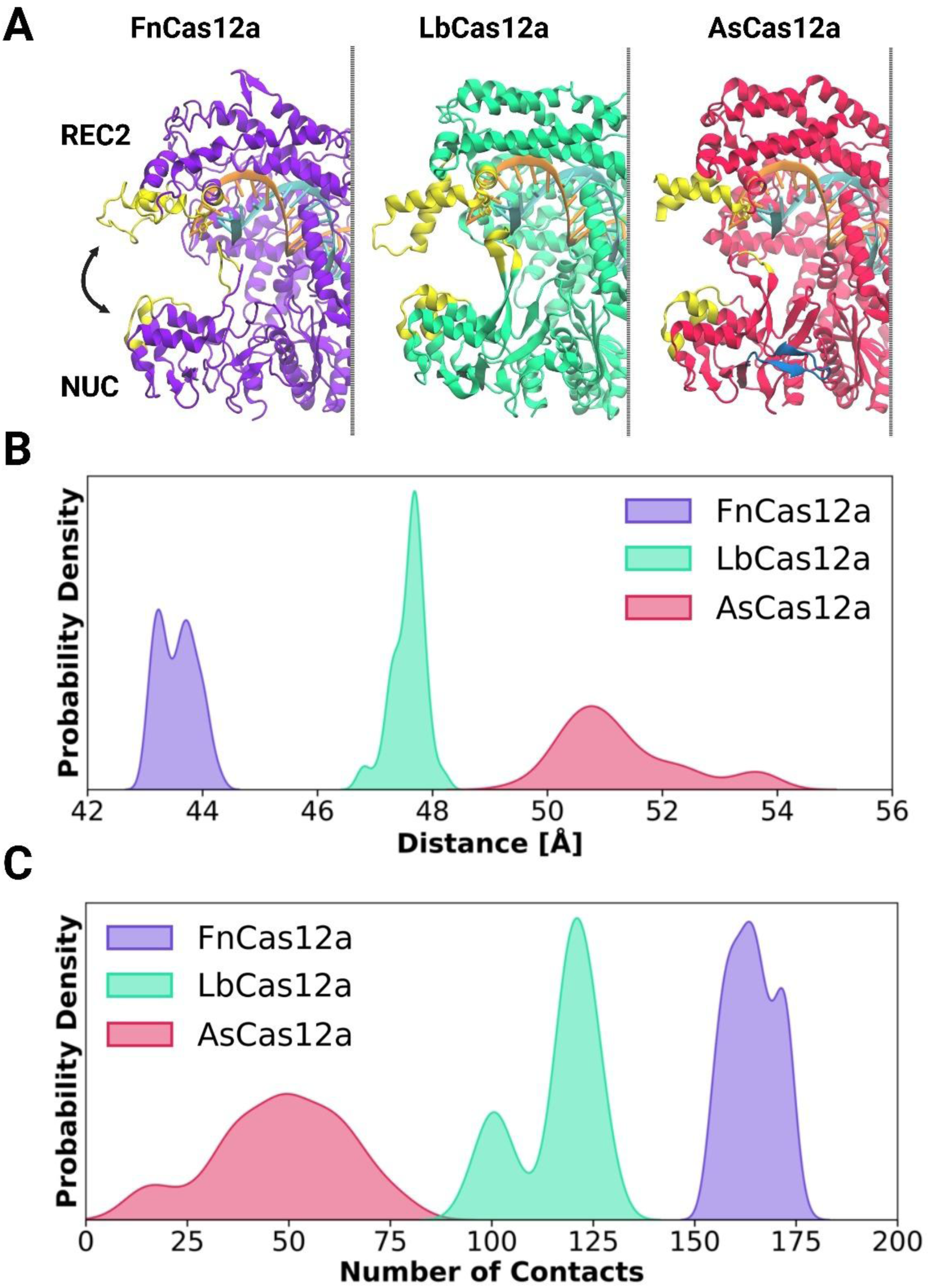
Molecular dynamics simulations of Cas12a ternary complexes. (**A**) Residues involved in REC2-NUC contacts highlighted in yellow, overlaying Fn Cas12a (6GTG, purple), Lb Cas12a (AF2 prediction, bright teal), As Cas12a (8SFR, red). The crRNA (orange) and target-strand (cyan) are shown, and the NUC loop 2 of AsCas12a (dark blue). Kernel density estimation plots of probability density for (**B**) distance between REC2 and NUC, and (**C**) probability density of contacts between REC2 and NUC.

These data show the three Cas12a orthologues have distinctly different conformational distributions. FnCas12a has probability density centred at 43-44 Å, LbCas12a at 47-48 Å, and AsCas12a has a broad distribution from 50-54 Å (**Figure 5B, Figure S19**). As the closed-conformation is thought to be particularly important for TS cleavage (24, 28, 30, 31), it is intriguing to note these distributions line up with observed rates of *k*_TS_ between Cas12a orthologues.

A cause, or consequence, of these distributions are the protein-protein contacts between the REC2 and NUC (**Figure 5C, Figure S20**). The probability of the number of contacts between the REC2 and NUC was quantified, and these show that FnCas12a has a high number of REC2-NUC contacts, peaking at ∼170 (**Figure 5C**). LbCas12a has two distinct peaks at ∼100 and ∼125 contacts, perhaps indicating two distinct conformations (**Figure 5C**). Strikingly, AsCas12a ranged from zero to almost one hundred contacts, peaking at ∼50 (**Figure 5C**).

In light of recent structural data characterising the conformation changes of the REC2 domain of AsCas12a in DNA cleavage (18), these REC2 – NUC dynamics are significant. Post NTS-cleavage, the REC2 of AsCas12a is highly flexible, and electron density was observed between the R-loop and NUC domain, suggesting the NUC loop is binding the crRNA:TS heteroduplex (18). By enumerating the residues involved in REC2 – NUC ‘pinching’ motions, we find an interplay between the NUC lobe, NUC loop, and REC2 domain.

## Discussion

We explored what drives the difference in function between Cas12a orthologues, using a combination of *in vitro* DNA cleavage assays, *in vivo* plasmid interference and genome editing, and *in silico* simulations. Our results show trade-offs between *cis* and *trans* cleavage, which may be driven by dynamic REC2 – NUC interactions.

### Kinetic comparison of wild-type Cas12a orthologues

We observed large differences in *cis* cleavage kinetics between Cas12a orthologues. These differences in strand cleavage kinetics are important, given a growing body of evidence that the kinetics of R-loop formation and DNA cleavage drive the target specificity of Cas12a nucleases (20, 65, 67, 68).

For LbCas12a and AsCas12a, the modelled values of *k*_NTS_ and *k*_TS_ were comparable to previously published values (18, 20, 25–27) (**Table S1**). The rates for FnCas12a were over 10x faster than a previous study (23), which was itself over 5x faster than another report (14) (**Table S1**). Notably, these studies both use half the Mg^2+^ ion concentration than herein (5 vs 10 mM MgCl_2_), which has been shown to decrease both DNA binding and *cis* cleavage rates for AsCas12a (20, 28). Furthermore, both previous kinetic studies for FnCas12a used short linear dsDNA substrates (14, 23). Previous work has shown negatively-supercoiled DNA substrates accelerate R-loop formation for LbCas12a, compared to the unconstrained topology of linear DNA substrates (27). Faster cleavage of plasmid DNA vs short oligonucleotides has also been observed for FnCas12a (23). The rapid plasmid DNA cleavage by FnCas12a in this study suggests it also has more rapid R-loop formation with negatively supercoiled DNA substrates, in contrast to AsCas12a, which has very similar *cis* cleavage kinetics between substrates (18, 20, 25) (**Table S1**).

The R-loop formation of AsCas12a has been studied in detail, and is thought to occur with minimal contribution from the REC domain (18, 20). The faster, and more torque sensitive, R-loop formation by FnCas12a and LbCas12a may indicate divergent Cas12a-heteroduplex interactions in target DNA recognition. Supporting this are high-throughput mismatch studies on plasmid DNA targets, which show that AsCas12a is much more specific than LbCas12a and FnCas12a (67). Critically, the kinetics of strand cleavage rates influence targeting specificity in a biological setting (69). Our comparison of wild-type FnCas12a, LbCas12a, and AsCas12a provide a kinetic explanation for these observed differences in specificity.

### REC2 and NUC loop interactions modulate strand cleavage kinetics

We replicated and expanded on previous work studying REC2 mutations (15, 26). We observed that alanine substitution of the REC2 ‘gate’ resulted in consistent increases in *k*_TS_ across Cas12a orthologues, to the detriment of *k*_NTS_ (**Figure 2A-C**). With rapid TS cleaving REC2 mutants, we observed slower *trans* cleavage when activated by full-length vs truncated TS (**Figure 2D-G**). This suggests TS-loading motions can sterically hinder *trans* ssDNA substrates. We propose the rapid TS-loading of FnCas12a contributes to its slow *trans* cleavage.

However, LbWT showed faster *trans* cleavage than its REC mutant LbW355A, for all combinations of crRNA and TS (**Figure 2D-G**). This would suggest factors other than steric-hindrance by the TS can reduce *trans* cleavage rates. Case in point, AsCas12a has very slow *k*_TS_, and *trans* cleavage activity as slow as FnCas12a (**Figure 2D-G**). We suggest that REC2 domain flexibilty is the cause. Flexibility of this domain results in RuvC lid ‘closing’ between NTS and TS cleavage (18). It is probable that the REC2 remains flexible post TS-cleavage, and this also reduces allosteric opening of the RuvC lid. Removing the REC2 stacking interaction likely also decreases heteroduplex affinity to the REC2, affording greater flexibility. Thus, REC2 flexibility – for WT or REC2 mutant nucleases – could be a cause of low *trans* cleavage.

Given REC2 mutants have faster TS cleavage than WT, it has been questioned why this aromatic interaction is conserved across the Cas12a family (26). Although we observed no defect in thermostability or in *E. coli* plasmid interference by these mutants (**Figure S3**), others have observed decreased gene editing by AsW382A in HEK293T cells (15). We suggest that these mutants can lose activity through lessened R-loop stability.

When complexed with a 20nt spacer crRNA, both LbWT and LbW355A showed more frequent occupation of shorter R-loop states, states that can lead to reversible DNA unwinding (26). However, LbW355A showed shorter R-loop states also with wild-type crRNAs (26). Interestingly, in cleavage assays using a crRNA with a 20nt spacer, they showed LbW355A had incomplete plasmid target linearisation – unlike compared to LbWT or assays using a 24nt spacer (26). We suggest that R-loop collapse after NTS cleavage, and before TS cleavage, is the cause of this incomplete cleavage.

Incomplete cleavage of target DNA (i.e. NTS nicking only) has also been observed at mismatched target sites (67, 70). These studies use negatively supercoiled target plasmids (67, 70), a substrate topology that can allow rapid R-loop formation (27). Mismatches between crRNA and TS decrease R-loop stability, and can lead to dissociation of Cas12a (20). In target-dependant nicking, Cas12a appears to form stable enough ternary complexes to permit NTS cleavage, but dissociate before TS cleavage (67, 70). Thus, we propose that alanine substitution of the REC2 ‘gate’ results in lower R-loop stability.

The FnΔLoop mutant also showed incomplete *cis* cleavage, leaving ∼20% of the target plasmid in the nicked state (**Figure S5B**). This activity is not likely to be caused by non-specific nicking by excess nuclease present in the single-turnover conditions, as this mutant is minimally *trans-*active (**Figure 4A**). Unlike the deletion mutants of LbCas12a and AsCas12a, this mutant displayed wild-type levels of thermostability and plasmid interference in *E. coli* (**Figure 3B**, **4D**). Thus, we suggest the FnΔLoop has incomplete target DNA cleavage through R-loop collapse and target DNA dissociation, caused by loss of critical NUC loop interactions. That NUC loop mutants have disrupted R-loop stability is consistent with recent structures capturing the NTS and TS cleavage states of AsCas12a (18). This work shows the NUC loop making numerous interactions with the NTS and TS when they are coordinated in the RuvC active site, and even suggests the NUC loop remains bound to the heteroduplex in-between NTS and TS cleavage (18).

Similar to LbW355A, the LbFLX mutant increased *k*_TS_ and decreased *k*_NTS_ (**Figure 3F**). LbFLX had showed very different activity between *E. coli* plasmid interference and human cell line editing, with significantly reduced activity in the latter (**Figure 4D, E**). We suggest that FLX substitution in LbCas12a weakens critical NUC-loop target strand interactions, resulting in disrupted strand cleavage order (*k*_TS_ > *k*_NTS_) and lower editing activity in human cell lines.

This was not the case for AsCas12a, where AsFLX had a specific effect on *k*_TS_ only (**Figure 3G**). This mutant retained wild-type levels of acitivity in *E.coli* plasmid interference and in editing human cell lines (**Figure 4D, F**). We propose this mutation minimally disrupts the R-loop formation process of AsCas12a. Likewise, but more trivally, the AsP1153A mutant was not significantly different to wild-type AsCas12a.

LbΔLoop, AsΔLoop, and AsFLX-2 mutants had relatively robust *cis* and *trans* cleavage at 30 °C (**Figure 3F, G**, **Figure 4B, C**), and poor activity in *E. coli* and/or human cell line editing at 37 °C (**Figure 4E, F**). Any interpretation regarding the effect of NUC loop disruption on their function is confounded by their globally lower stability (**Figure 3C, D**).

### Molecular dynamics simulations reveal inter-orthologue differences in dynamic states

Molecular dynamics simulations showed the REC2 and NUC domain make numerous contacts in their dynamic motions, the REC2 ‘gate’ and NUC loop notably amongst them (**Figure 5A, Table S4**). This lends credence to the notion that the NUC loop aids the TS to traverse the distance between REC2 ‘gate’ and RuvC.

The predicted distance distributions, FnCas12a < LbCas12a < AsCas12a, fit well with observed TS cleavage rates (**Figure 5B**, **Figure 1H**). Given the importance of REC2 – NUC dynamics in TS cleavage, we propose this is a major driver of *k*_TS_. However, there remain large and unexplained differences in *k*_NTS_ between Cas12a orthologues.

The predicted REC2 - NUC domain contacts are particularly interesting in light of a study comparing the one-dimensional DNA diffusion of FnCas12a, LbCas12a, and AsCas12a (65). This work determined that FnCas12a has notably slower diffusion than LbCas12a and AsCas12a, and that an alpha helix in the REC2 plays a key role in this diffusion (65). Alanine substitution of the positively charged residues in this helix reduced DNA diffusion rates, and reduced genome editing efficiencies in HEK293T cells (65). In our MD simulations, these richly charged alpha helices are predicted to make dynamic contacts with the NUC (**Figure S21**).

This raises the possibility that this critical REC2 alpha-helix may be occluded by the dynamic pinching motions of Cas12a – thus limiting DNA diffusion and genome editing. This rests on the assumption that observed REC2 – NUC dynamics (Fn < Lb < As) also occur in the binary state. Evidence in favour of this notion are binary state structures of FnCas12a and LbCas12a (4, 17), which recapitulate REC2-NUC contacts predicted in MD simulations, with the exception of NUC loop contacts (**Figure 5A**, **Table S4**). Furthermore, diffusing smFRET experiments on apo and binary FnCas12a show free transitions between open and closed states, with 70 to 80% of molecules adopting the closed conformation (22).

**Figure 6:**
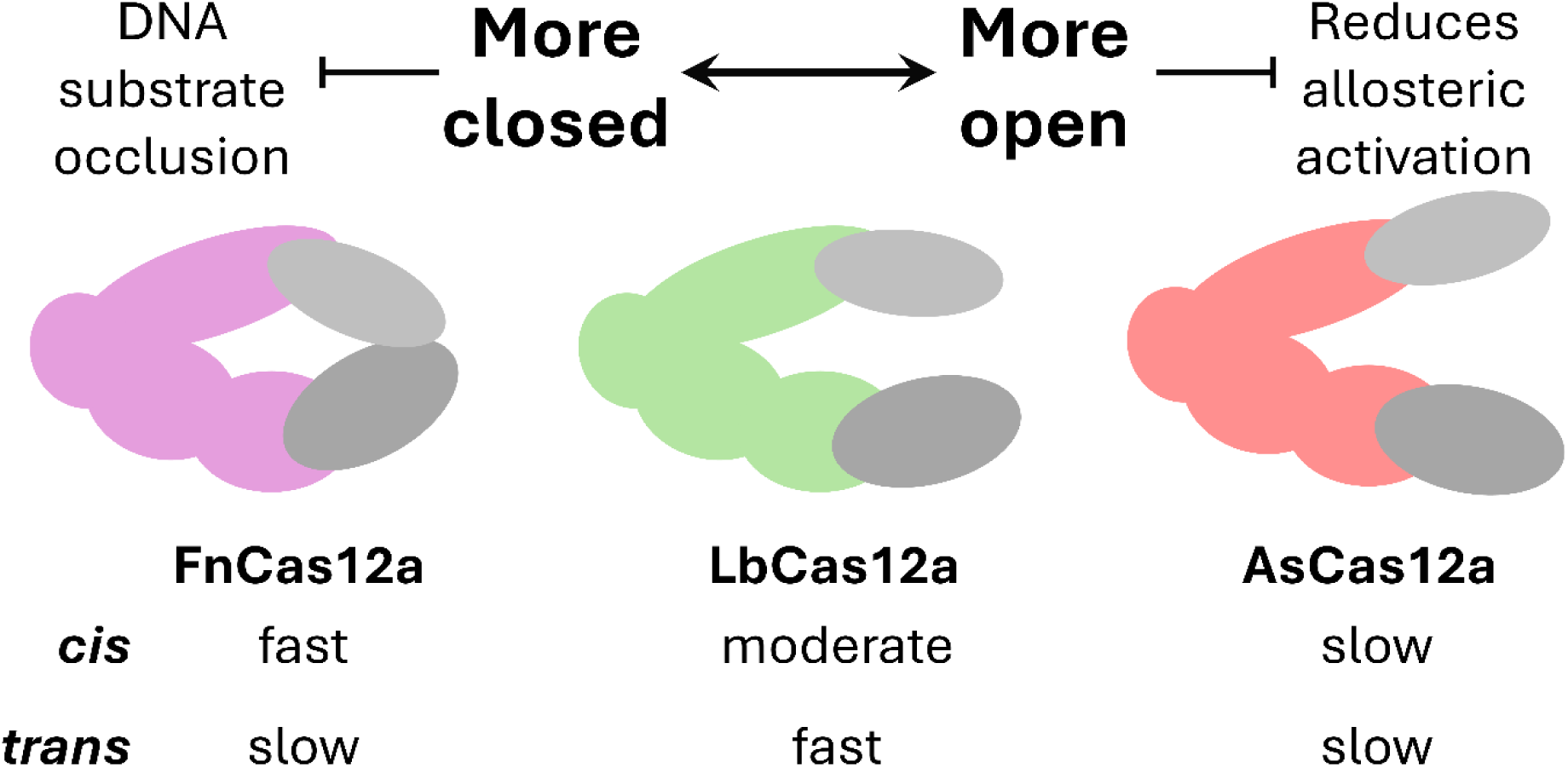
A model demonstrating trade-offs between cis and trans cleavage, driven by REC2-NUC interactions.

## Conclusions

In total, this comparative study of three Cas12a orthologues highlights trade-offs between *cis* and *trans* cleavage, driven by REC2 and NUC loop interactions. We highlight the critical role of the NUC loop in modulating DNA cleavage kinetics of Cas12a orthologues. This survey of structural and dynamic differences between Cas12a orthologues provides a mechanistic basis for their different abilities in genome editing and DNA detection, and lays the groundwork for future engineering of Cas12a nucleases.

## Data availability

Datasets generated herein are available on request.

## Acknowledgements

The authors would like to thank the Biomolecular Resource facility at the ANU for undertaking the DNA sequencing.

## Author contributions

A.N. performed plasmid cloning and protein purification, and assays for protein thermostability, DNA *cis* and *trans* cleavage, and *E. coli* plasmid interference. L.S. performed human cell culturing and editing. A.N. and G.B. performed kinetic modelling of *cis* cleavage rates. A.N. and G.B. analysed the high-throughput sequencing data. A.S. and P.R.A. performed and analysed the molecular dynamics simulations. A.S. contributed to writing the computational results and methods sections of the manuscript. G.P. conceived the computational investigations and supervised A.S.’s writing. A.N. and G.B conceived the study and wrote the manuscript. All authors commented and approved the manuscript.

