## Supplementary Files for "CRISPR-Cas12a REC2 – NUC interactions drive target-strand cleavage and constrain trans cleavage"

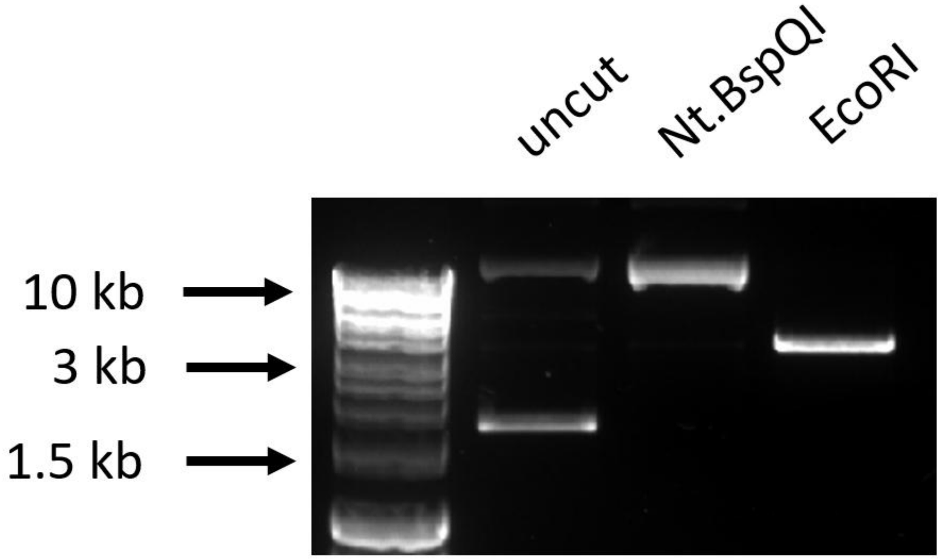


**Figure S1:** *Changes in target plasmid topology with nicking by Nt.BspQI, and linearisation with EcoRI.*

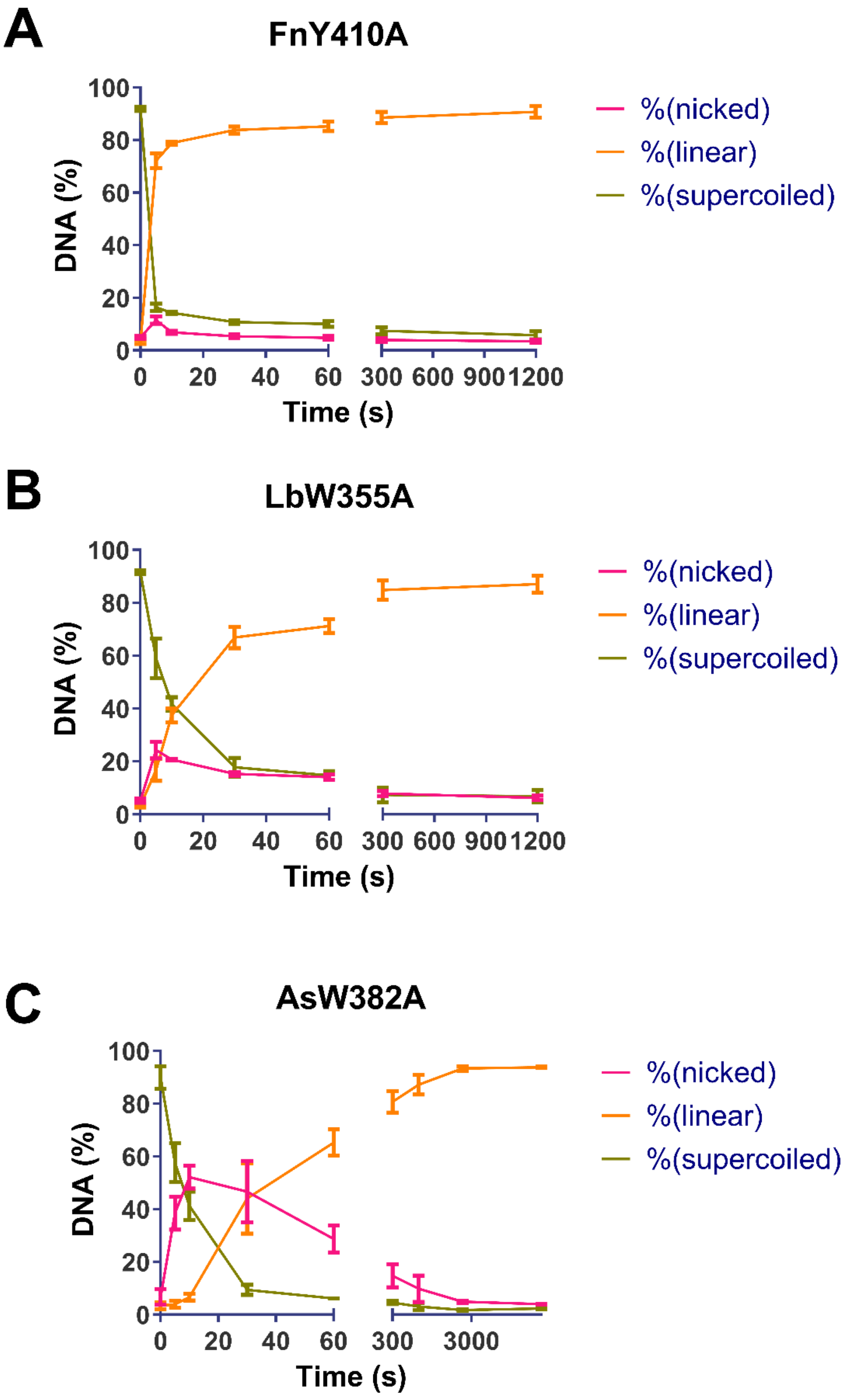


**Figure S2:** *Quantification of DNA fractions over time, with Cas12a mutant indicated. Line shows mean, error bars s.d.*

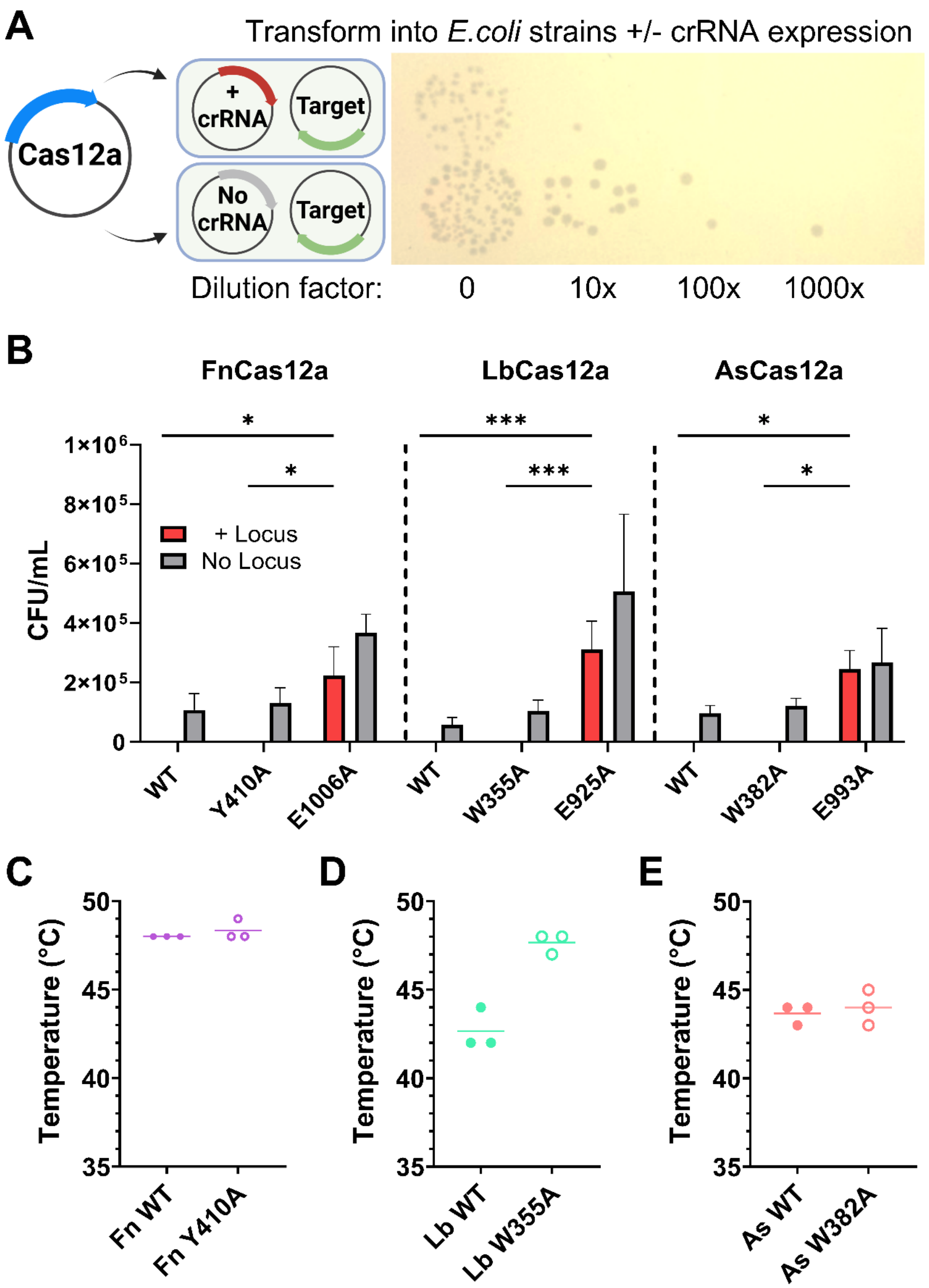


**Figure S3:** *(A) Outline of plasmid interference assay, image shows example transformation. (B) Mean colony forming units per mL (error bars show s.d.), for +/- crRNA conditions, Cas12a as indicated. Statistical significance evaluated by two-way ANOVA with Tukey's multiple comparison test (*p<0.1, **p<0.01, ***p<0.001). (C) Thermostability assay, melting point defined as fluorescence peak, line shows mean of three replicates.*

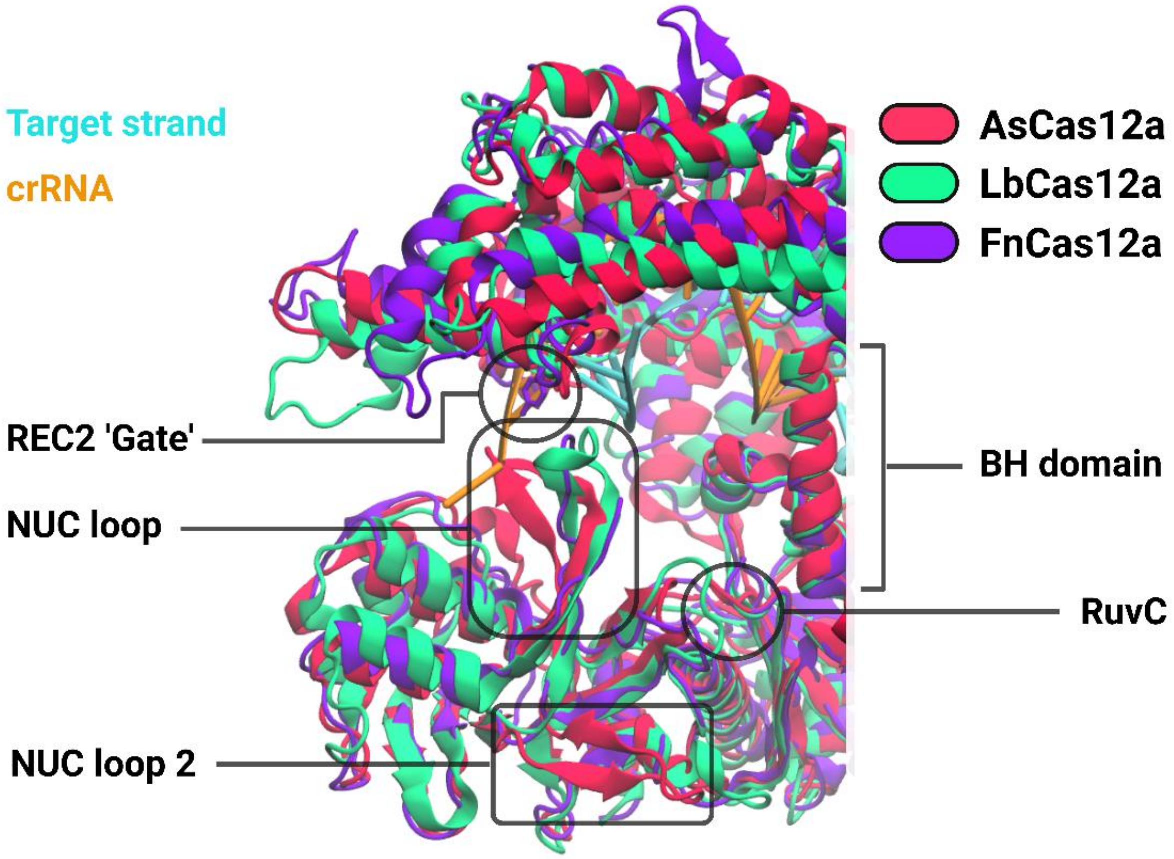


**Figure S4:** *Structural alignment of Cas12a orthologues, highlighting key features. AsCas12a (AF2 model, red), LbCas12a (AF2 model, bright teal), and FnCas12a (6GTG, purple).*

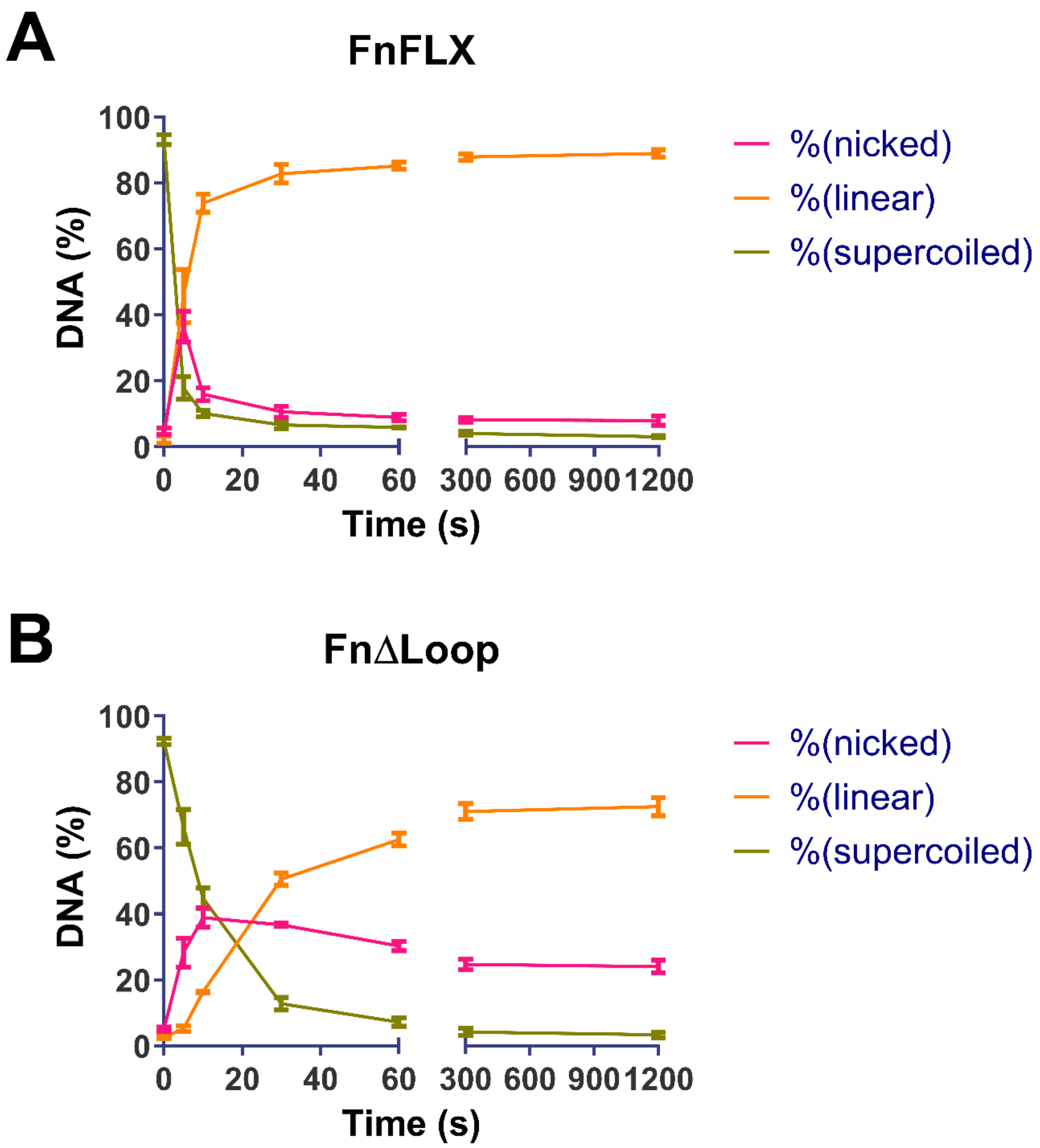


**Figure S5:** *Quantification of DNA fractions over time, with Cas12a mutant indicated. Line shows mean, error bars s.d.*

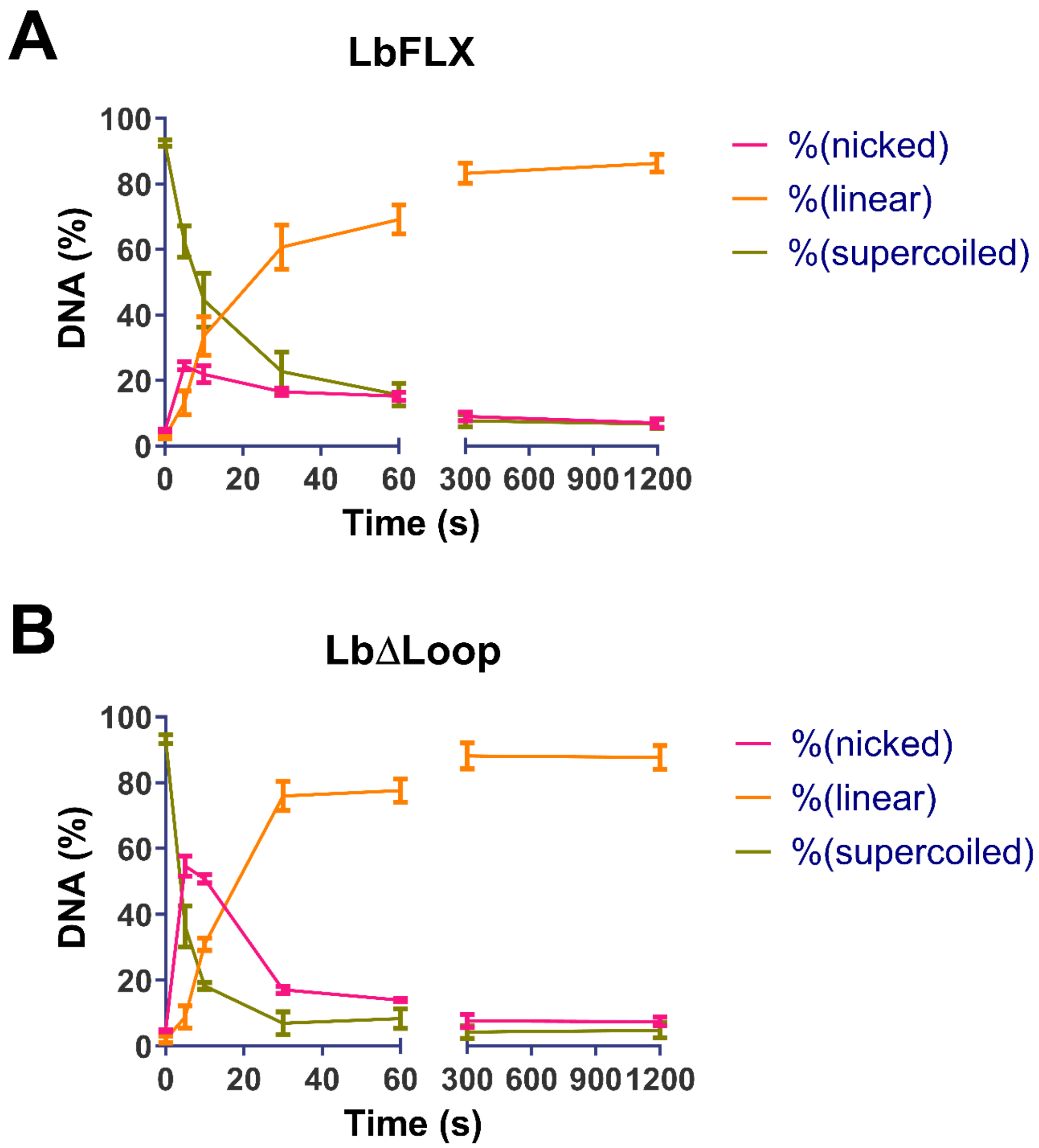


**Figure S6:** *Quantification of DNA fractions over time, with Cas12a mutant indicated. Line shows mean, error bars s.d.*

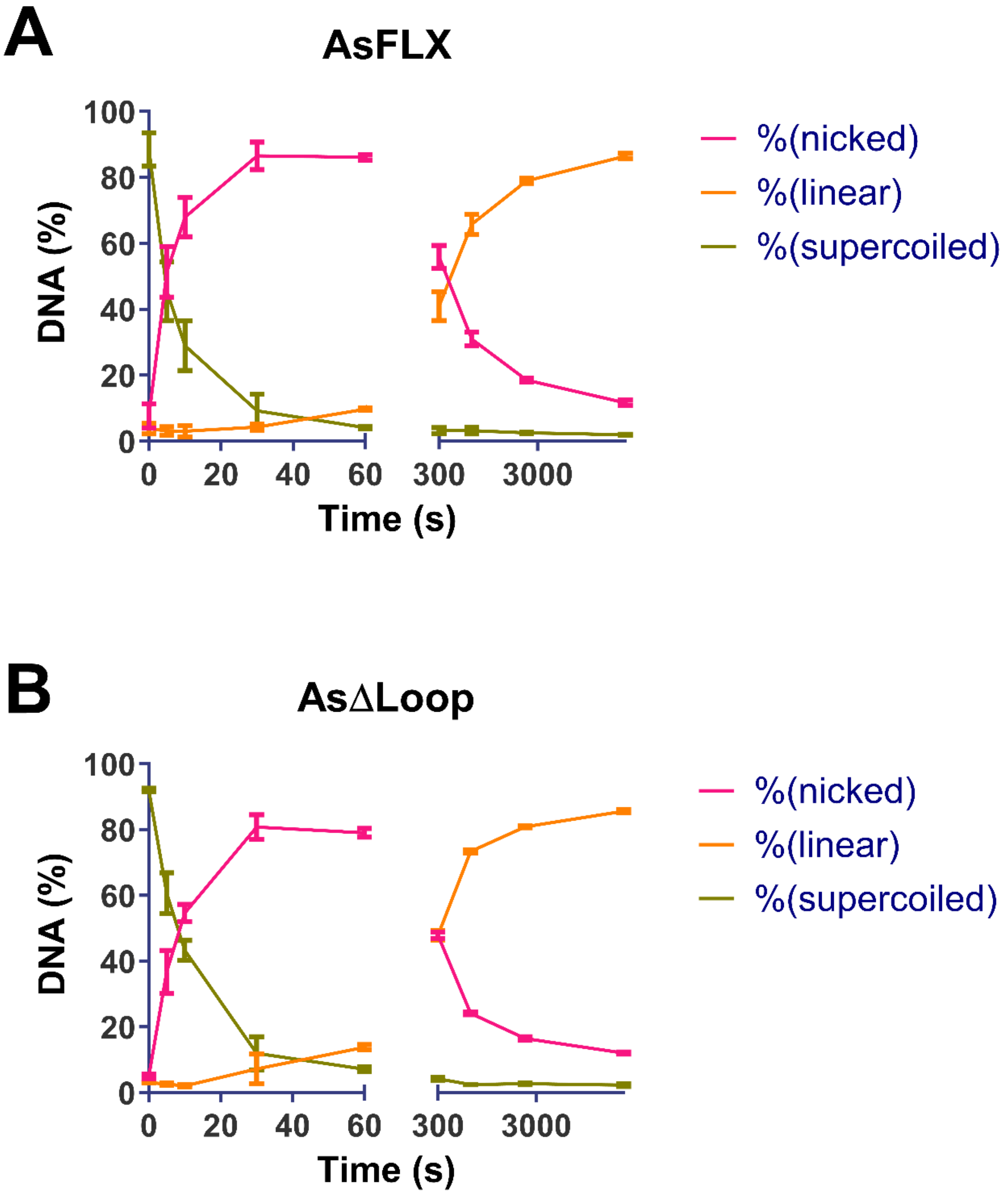


**Figure S7:** *Quantification of DNA fractions over time, with Cas12a mutant indicated. Line shows mean, error bars s.d.*

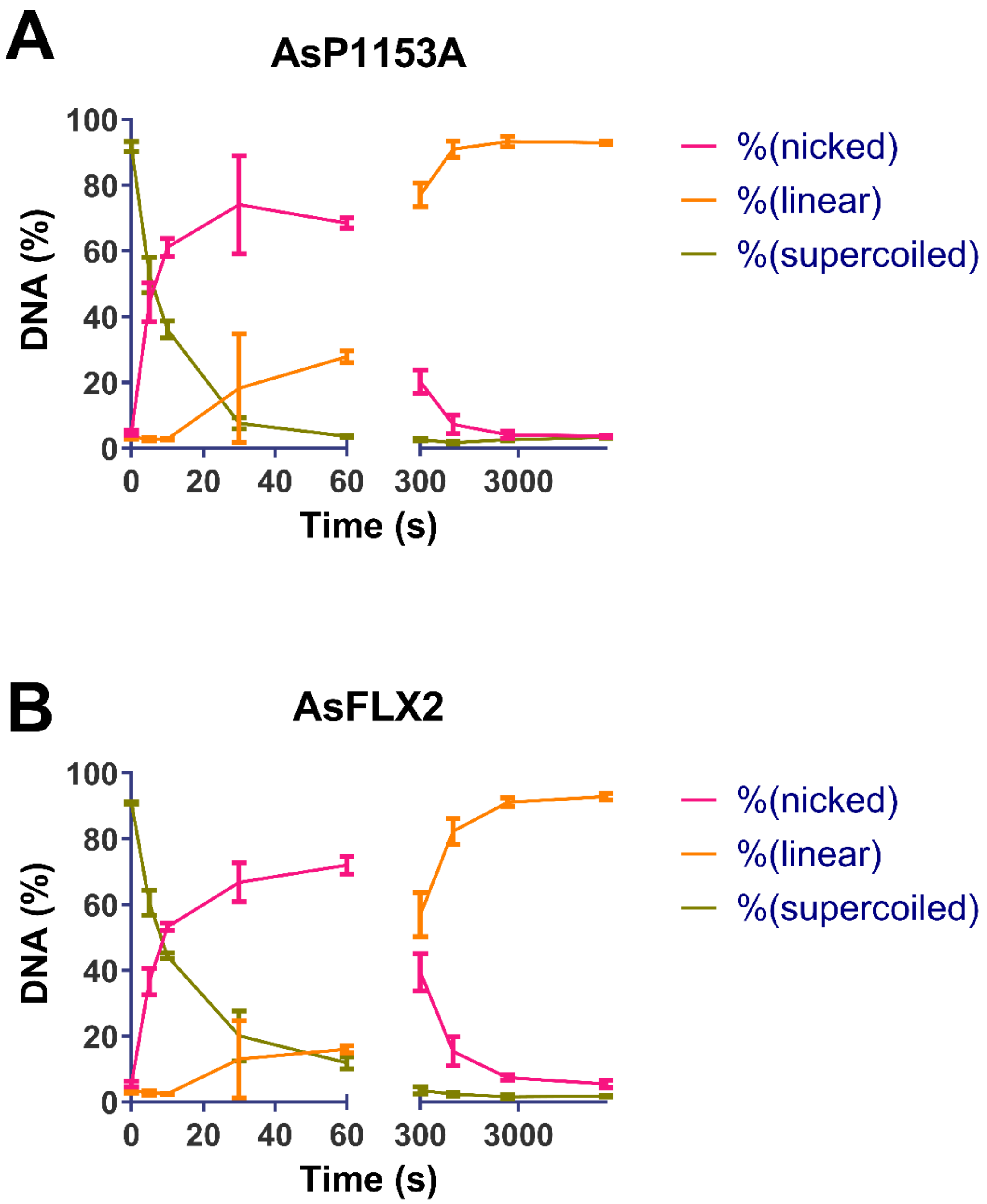


**Figure S8:** *Quantification of DNA fractions over time, with Cas12a mutant indicated. Line shows mean, error bars s.d.*

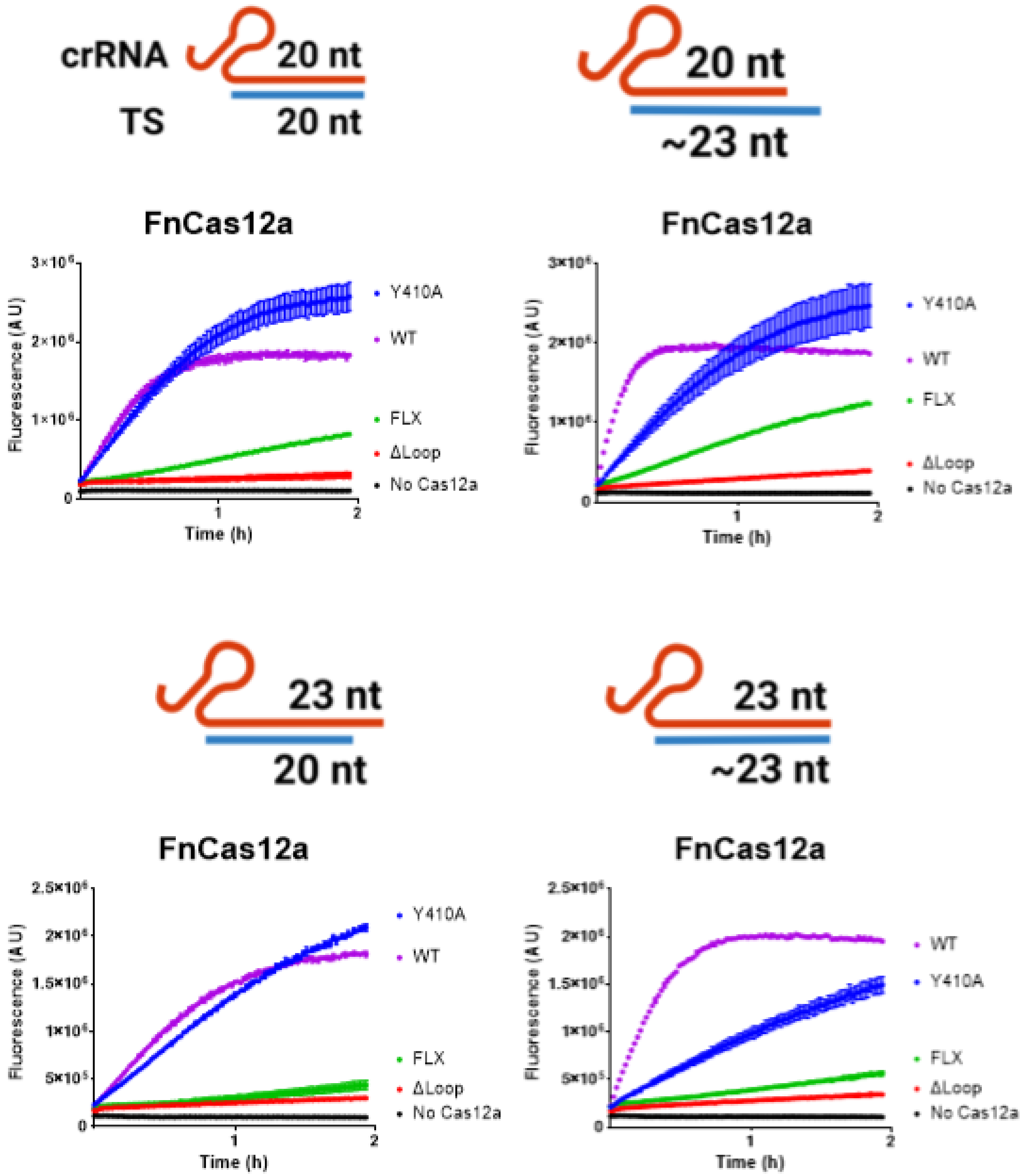


**Figure S9:** *Trans cleavage curves, with crRNA and TS combination indicated. Points show mean, error bars s.d.*

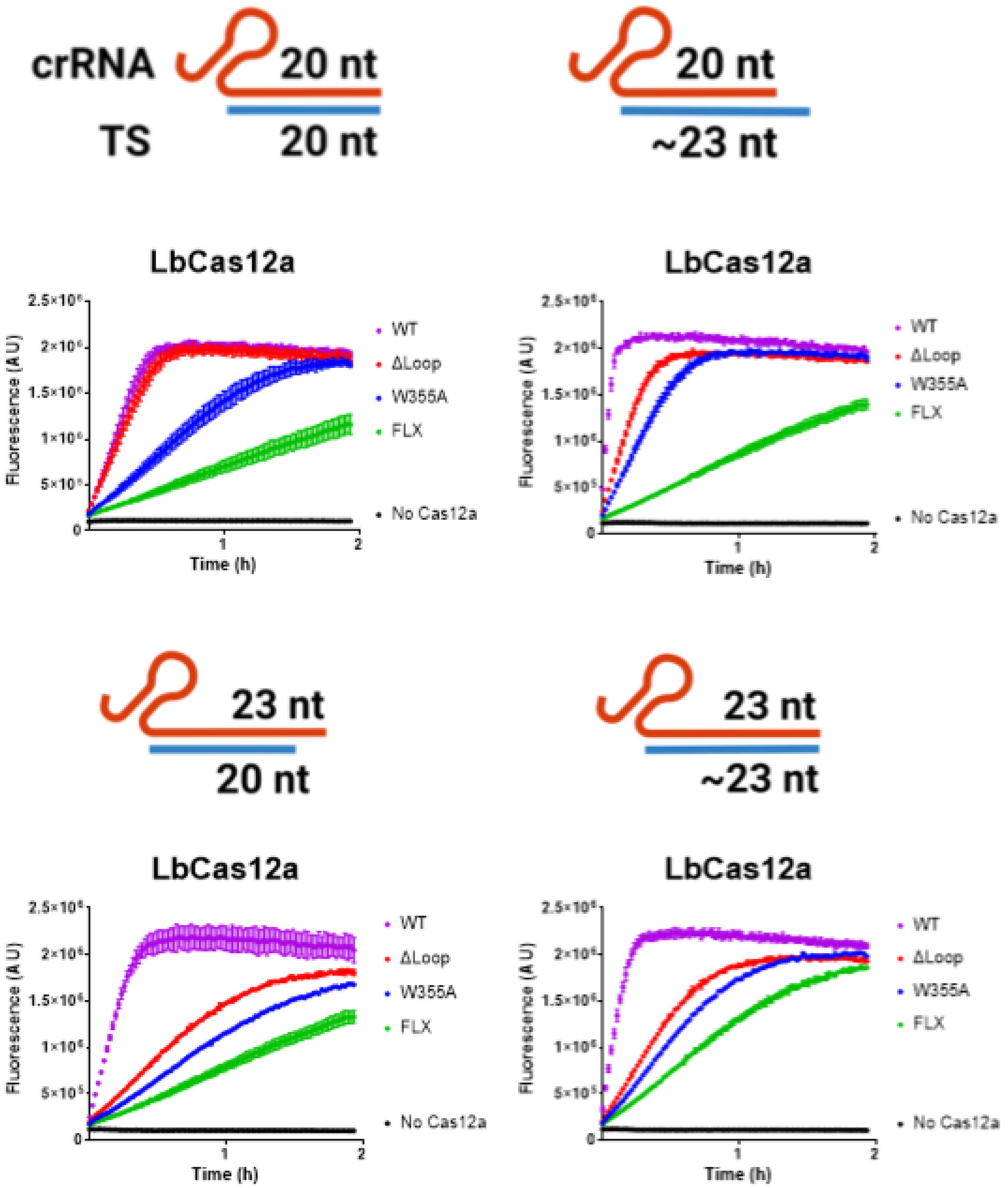


**Figure S10:** *Trans cleavage curves, with crRNA and TS combination indicated. Points show mean, error bars s.d.*

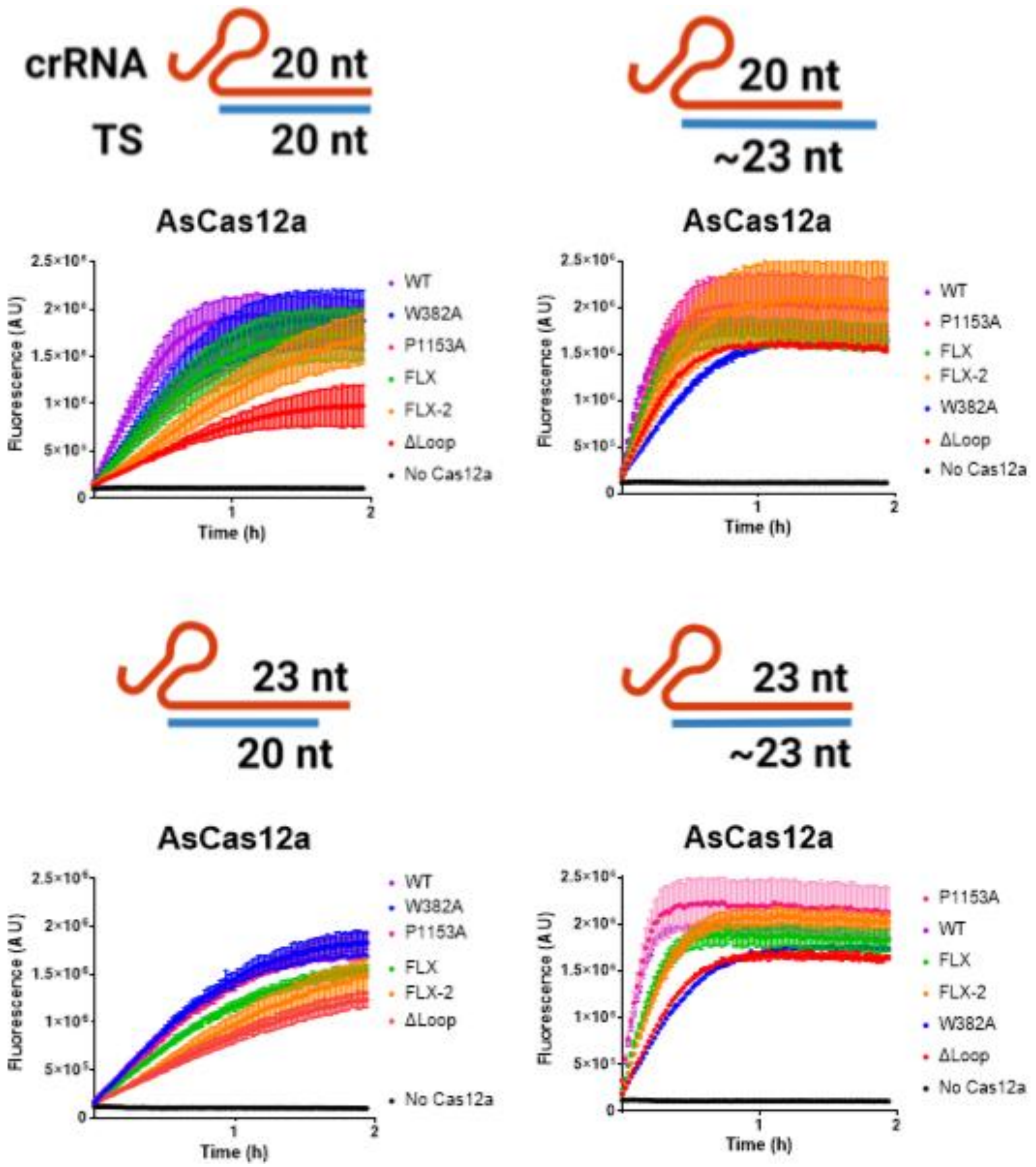


**Figure S11:** *Trans cleavage curves, with crRNA and TS combination indicated. Points show mean, error bars s.d.*

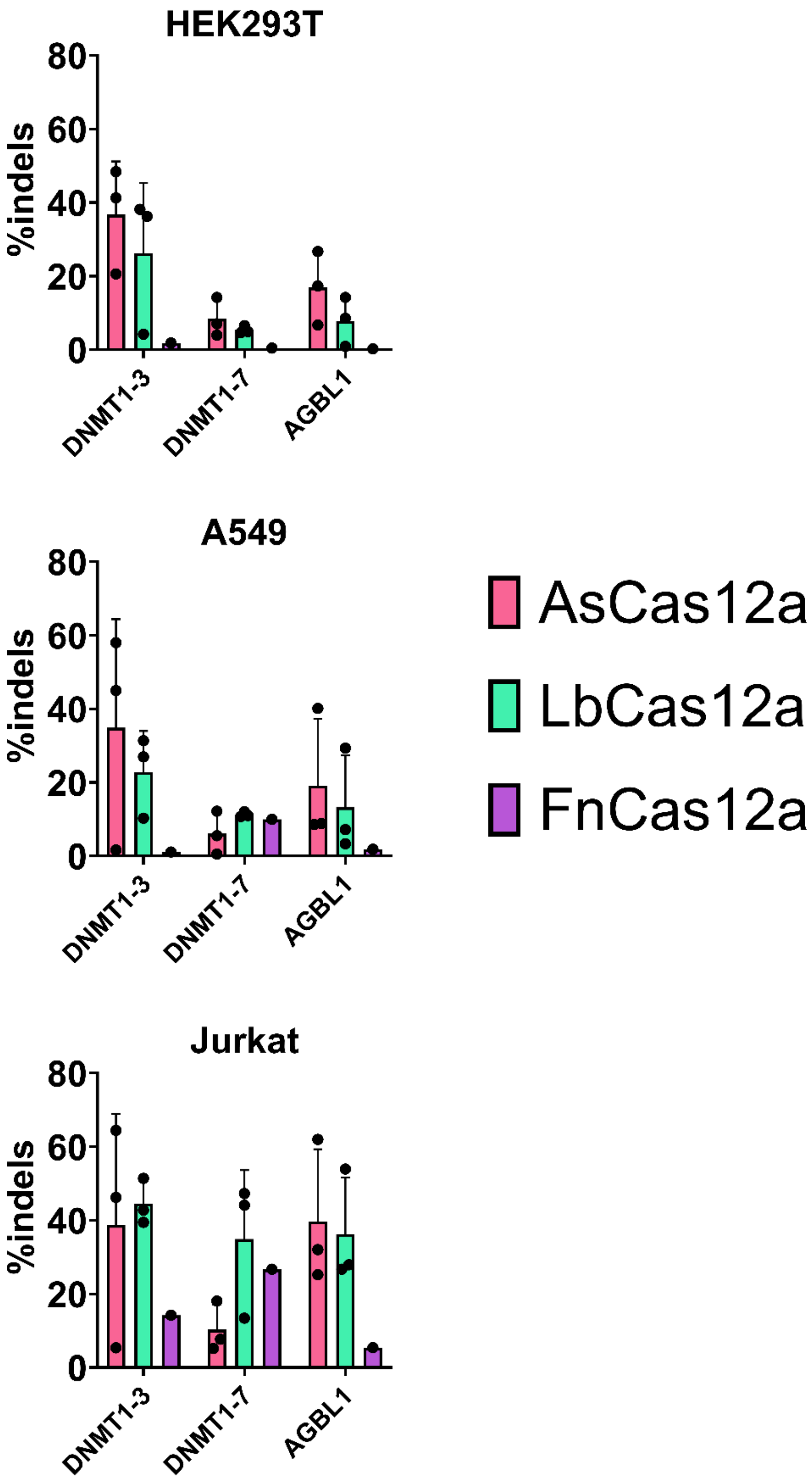


**Figure S12:** *Gene editing in cell line indicated, with crRNA indicated. Dots shown individual values, bar shows mean, error bars s.d. Note, n = 1 transformations performed for FnCas12a.*

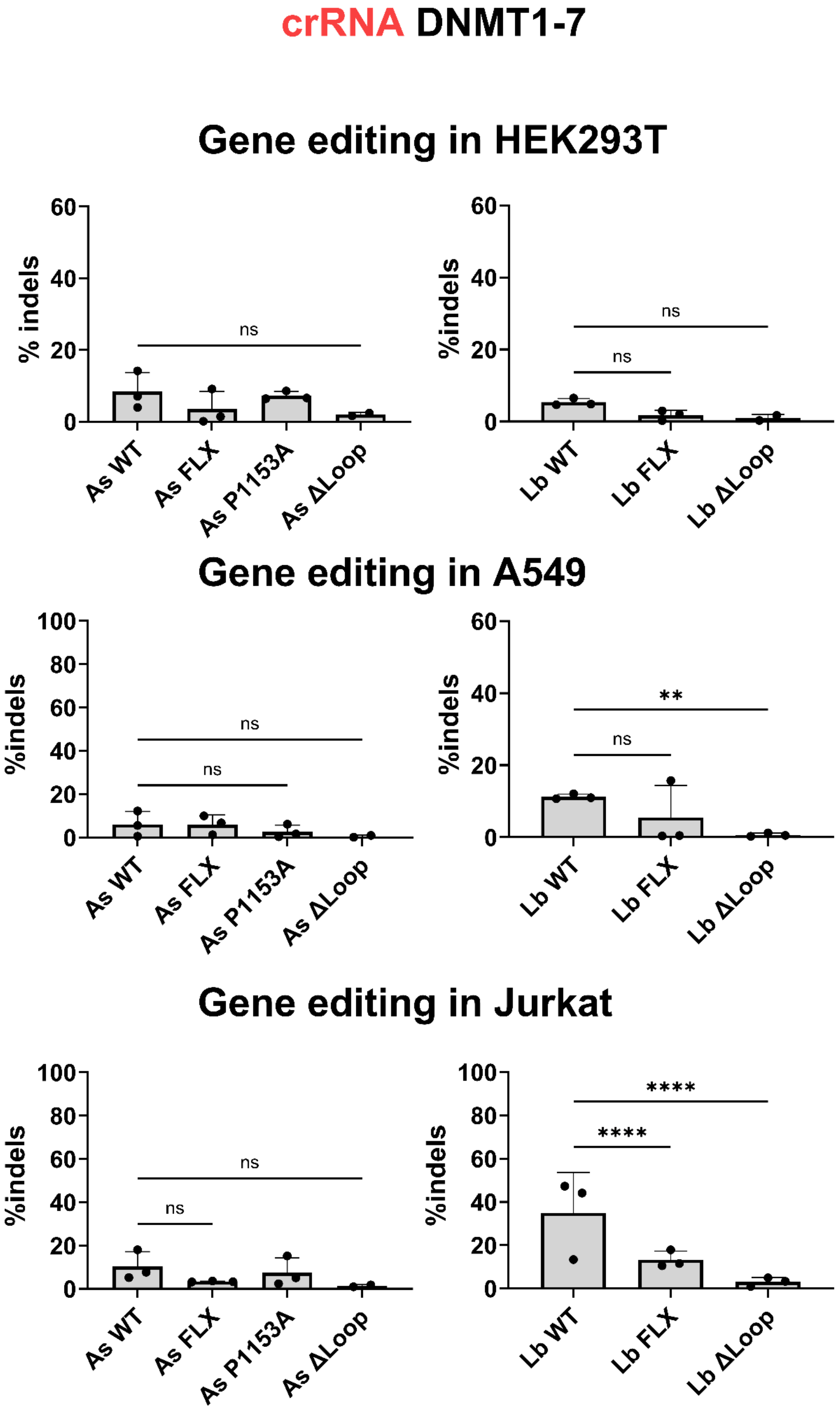


**Figure S13:** *Gene editing in cell line indicated, with crRNA DNMT1-7. Bars show mean, error bars s.d. Statistical significance evaluated by two-way ANOVA with Tukey's multiple comparison test (*p<0.1, **p<0.01, ***p<0.001, ****p<0.0001).*

### crRNA DNMT1-3

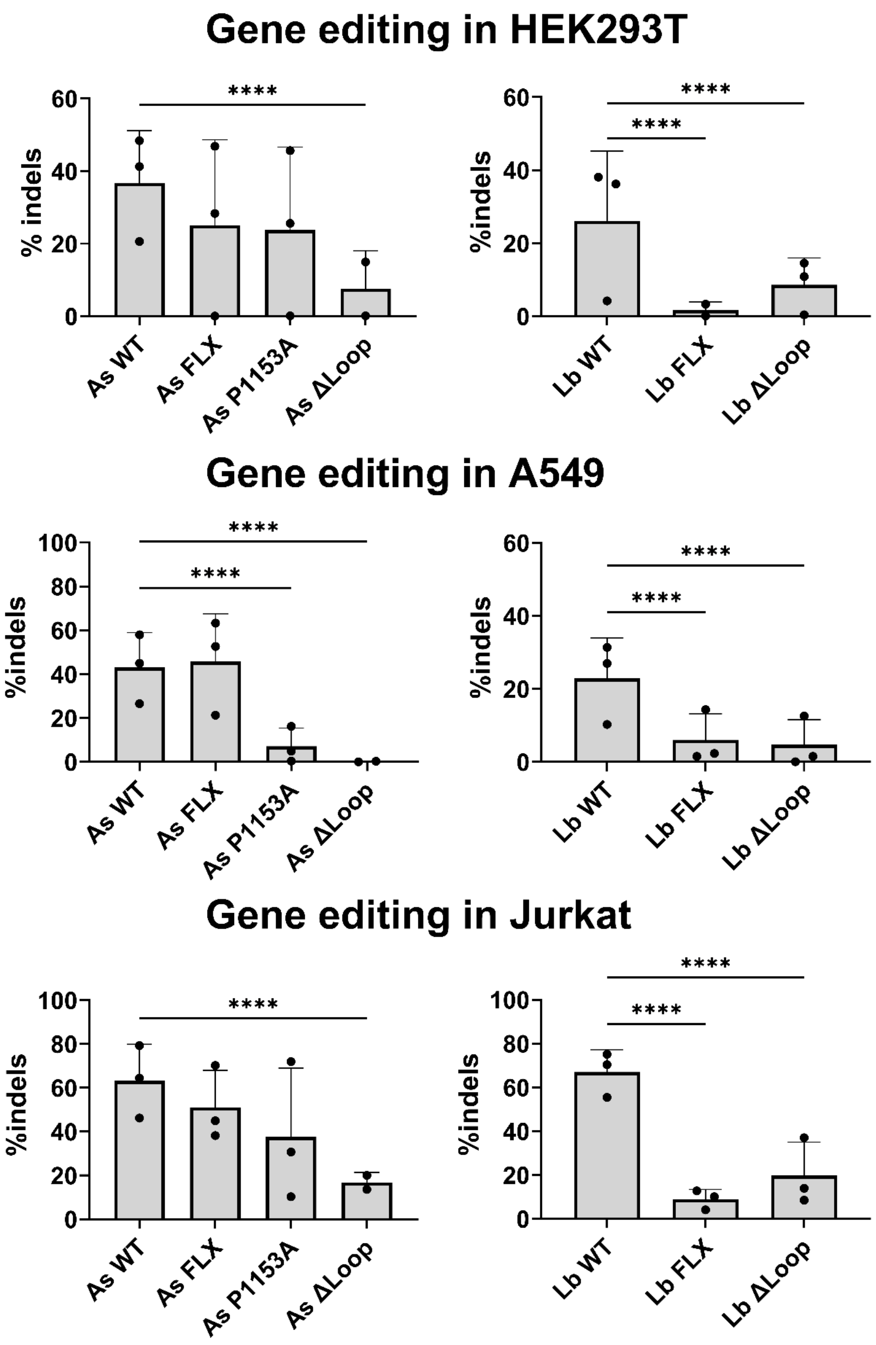


**Figure S14:** *Gene editing in cell line indicated, with crRNA DNMT1-3. Bars show mean, error bars s.d. Statistical significance evaluated by two-way ANOVA with Tukey's multiple comparison test (*p<0.1, **p<0.01, ***p<0.001, ****p<0.0001).*

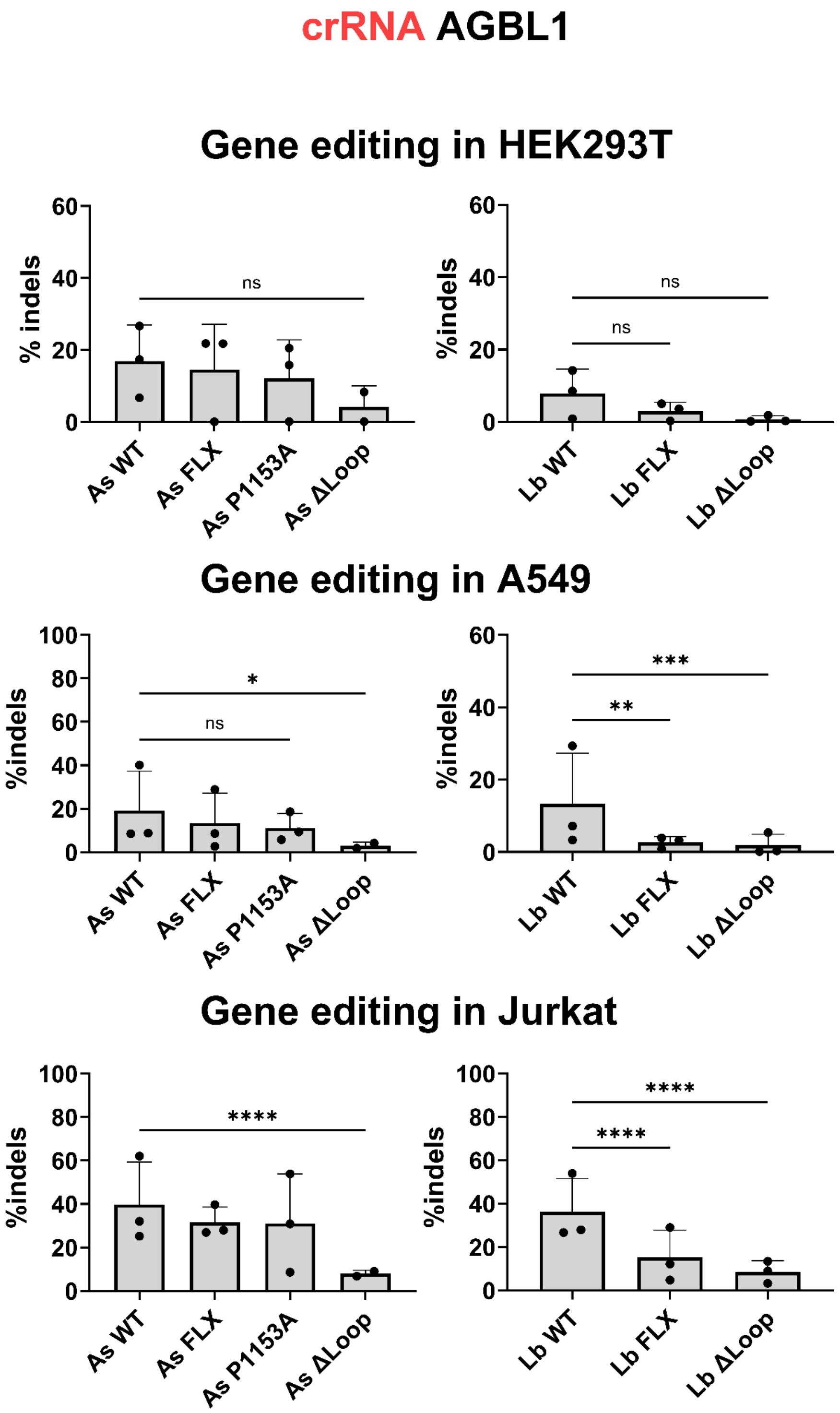


**Figure S15:** *Gene editing in cell line indicated, with crRNA AGBL1. Bars show mean, error bars s.d. Statistical significance evaluated by two-way ANOVA with Tukey's multiple comparison test (*p<0.1, **p<0.01, ***p<0.001, ****p<0.0001).*

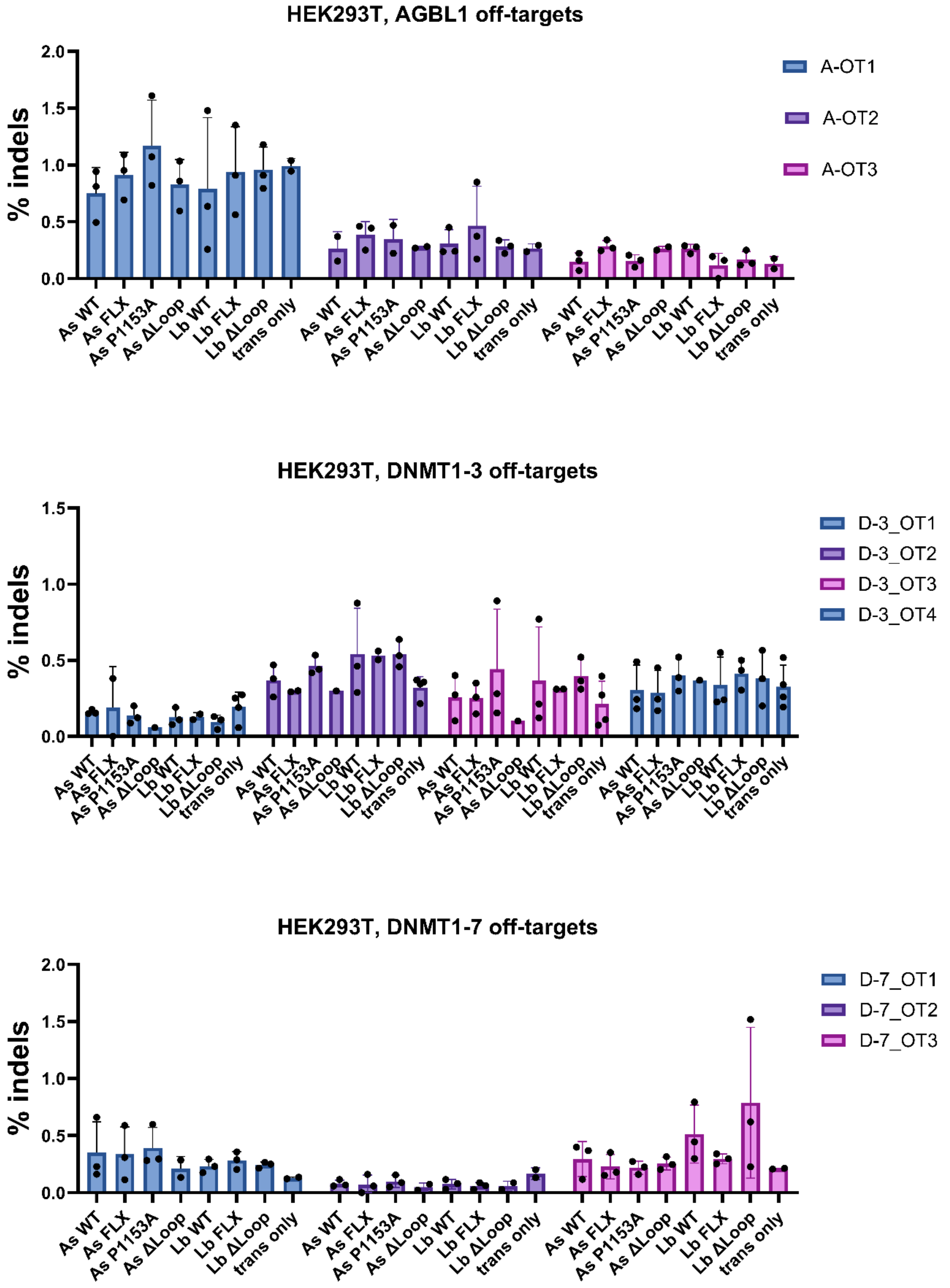


**Figure S16:** *Off-target (OT) editing with crRNA indicated, in HEK293T cell line. Bars shown mean, error bars s.d.*

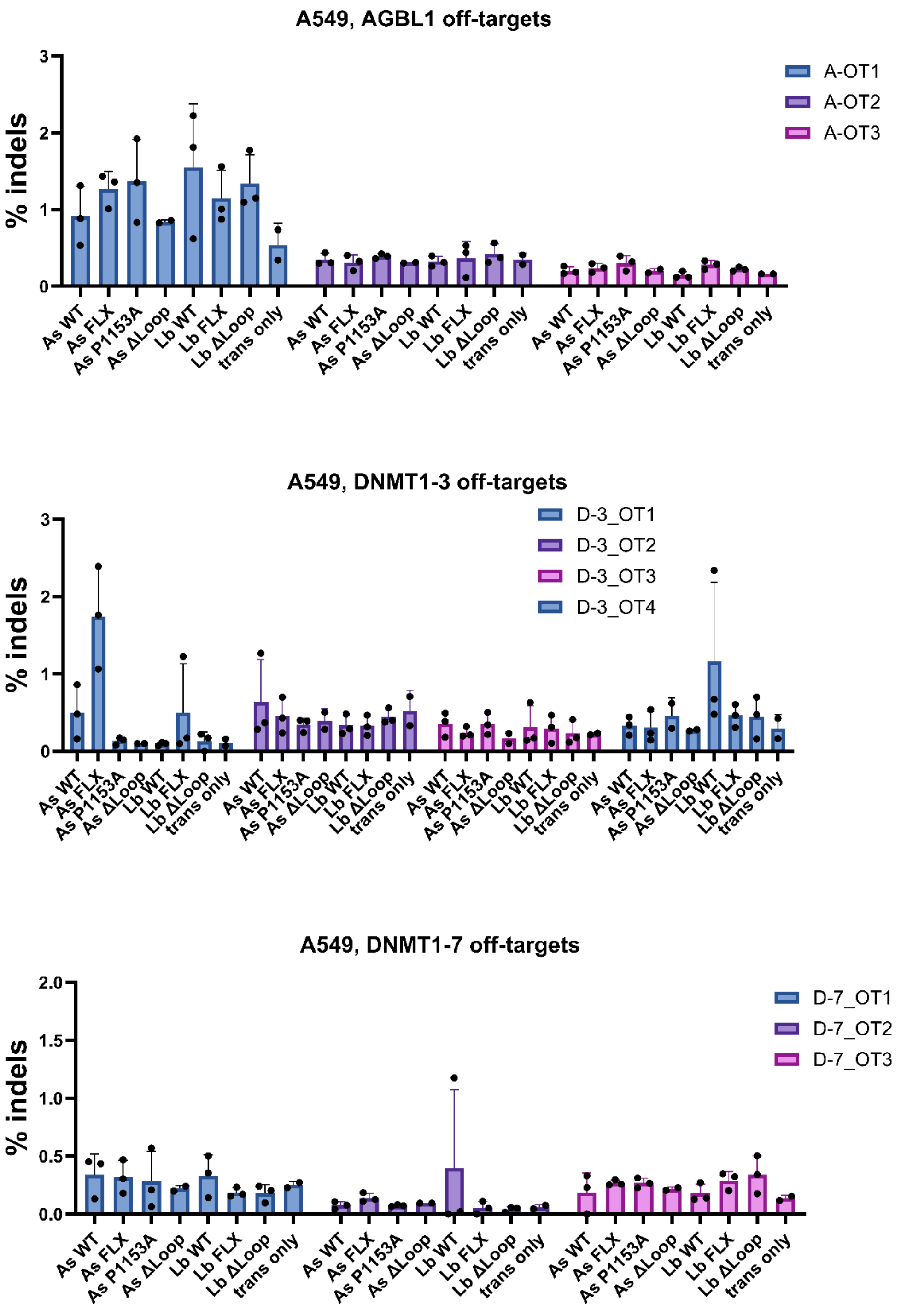


**Figure S17:** *Off-target (OT) editing with crRNA indicated, in A549 cell line. Bars shown mean, error bars s.d.*

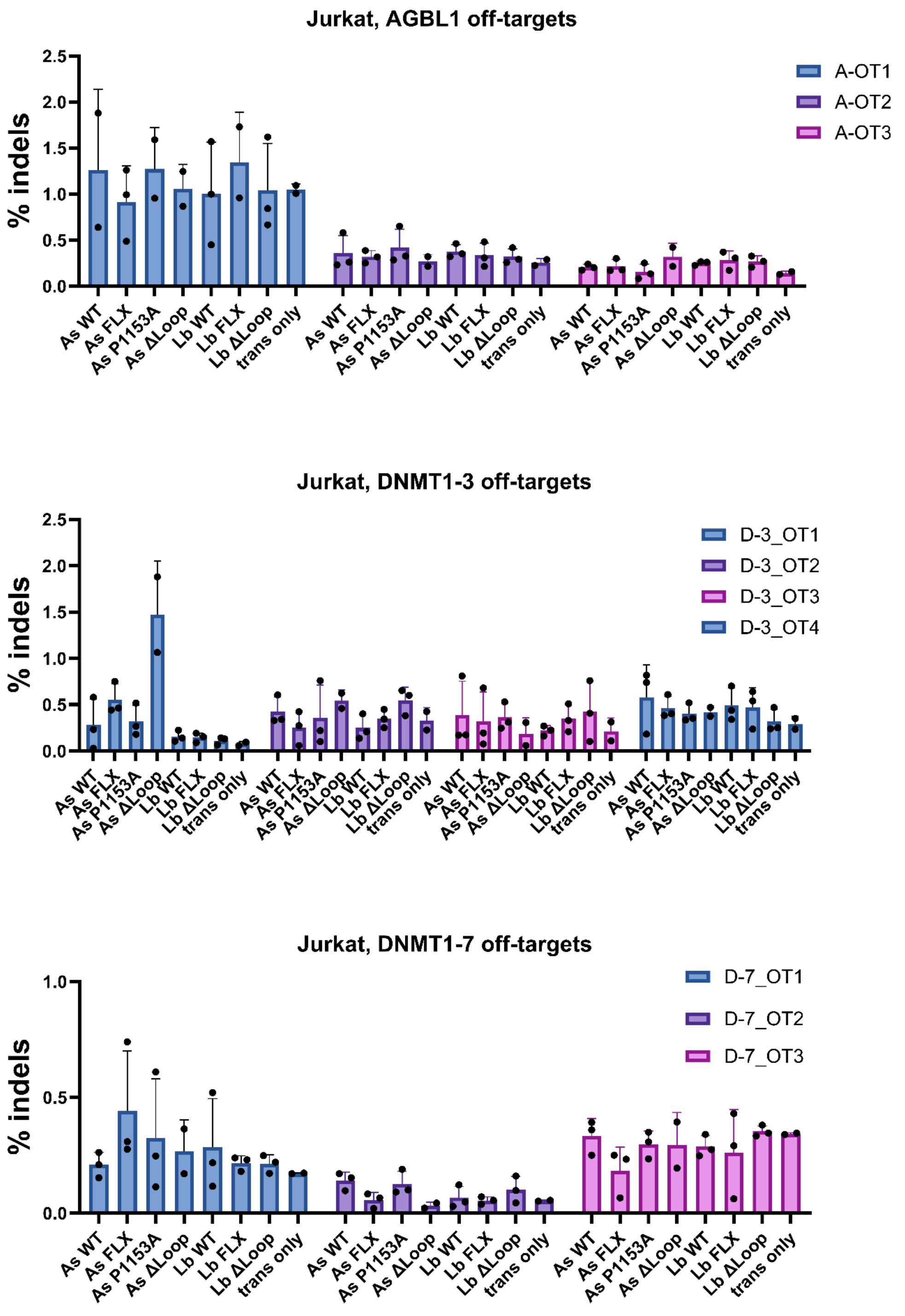


**Figure S18:** *Off-target (OT) editing with crRNA indicated, in Jurkat cell line. Bars shown mean, error bars s.d.*

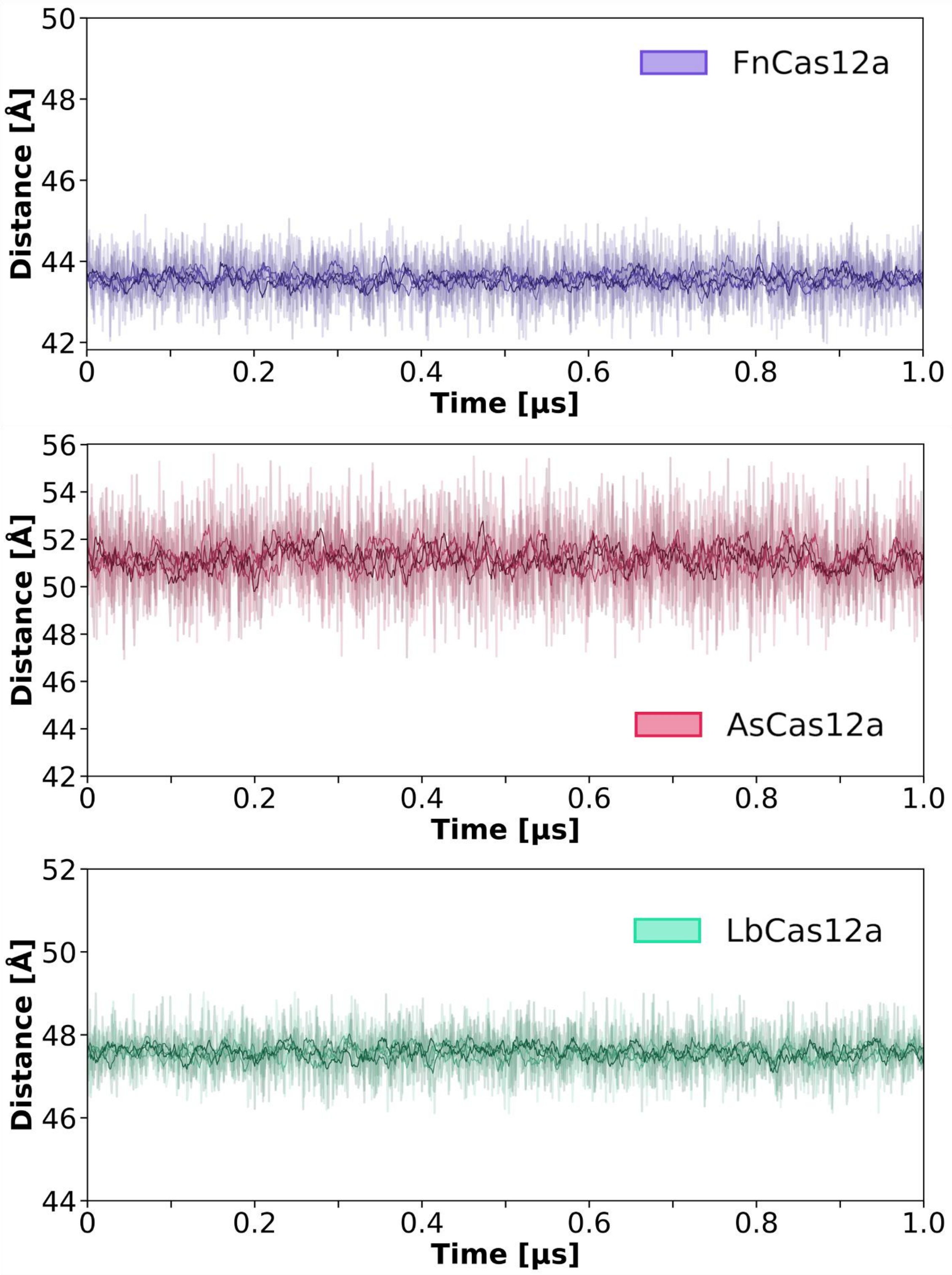


**Figure S19:** *Distance analysis of replicates of μs – length simulations.*

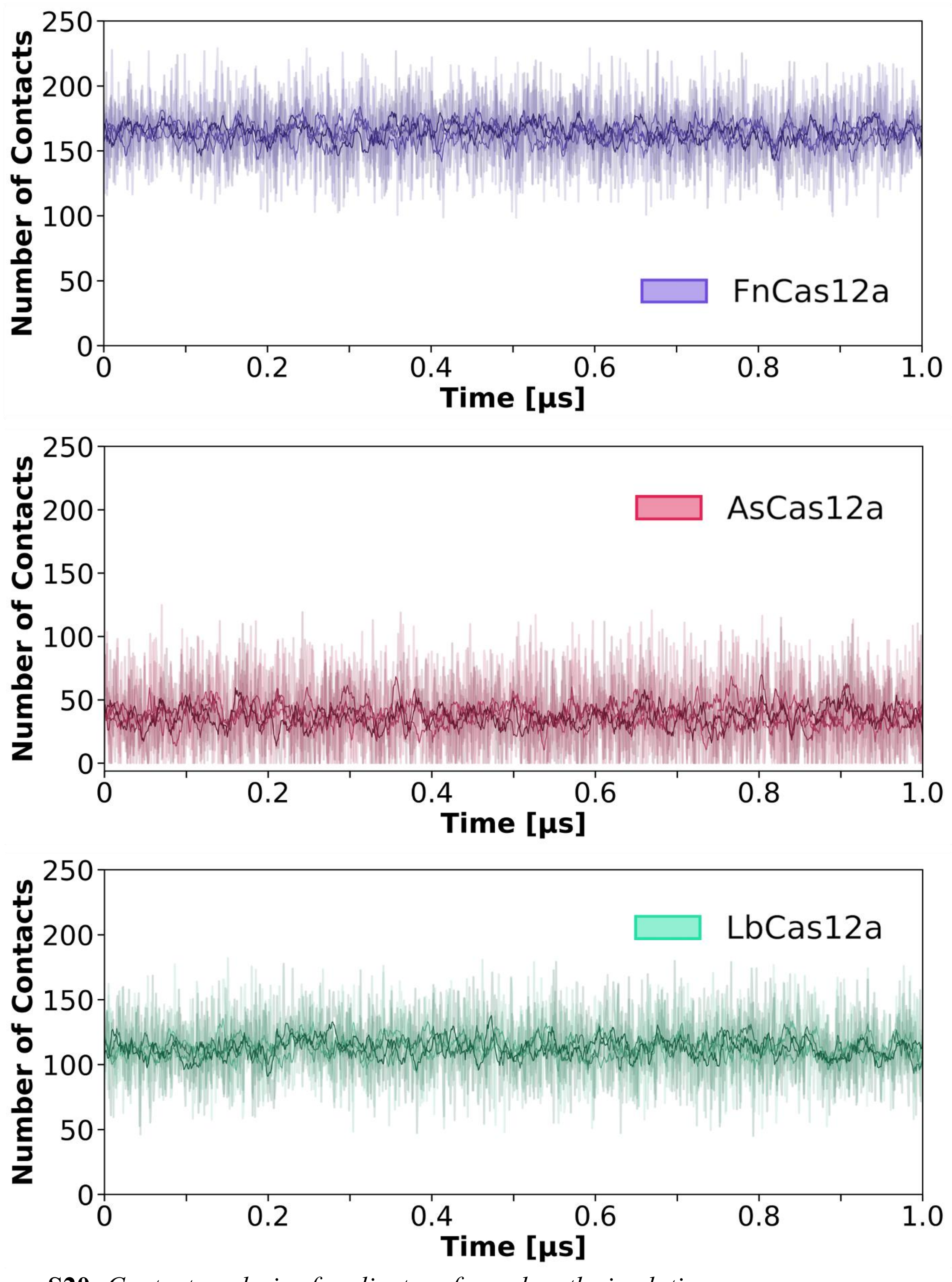


**Figure S20:** *Contact analysis of replicates of μs – length simulations.*

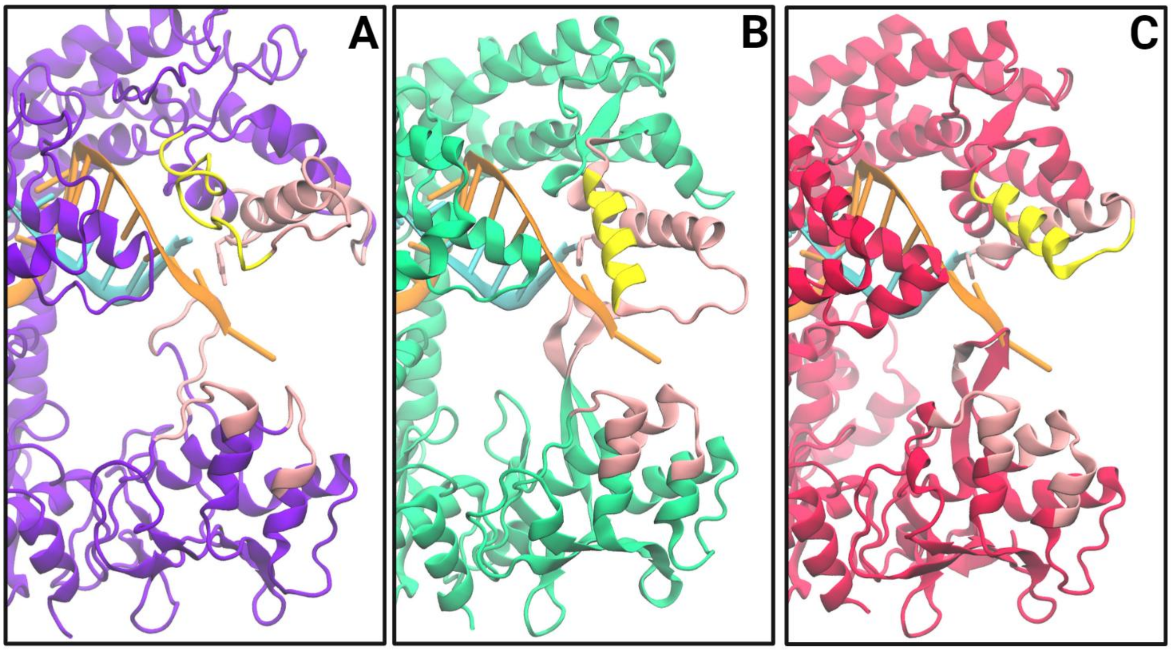


**Figure S21:** *Schematic of residues involved in REC2-NUC interactions (pink), and 1D DNA diffusion (yellow), overlaying (A) FnCas12a -6GTG, (B) LbCas12a -AF2 model, and (C) AsCas12a -AF2 model.*

### Supplementary Tables

| Cas12a | T (°C) | $k_{NTS}$ ($s^{-1}$) | $k_{TS}$ ($s^{-1}$) | DNA topology | Buffer |
|---|---|---|---|---|---|
| Lb[27] | 25 | 0.203 ± 0.048 | 0.012 ± 0.004 | (-) SC plasmid | 10 mM Tris-HCl, pH 7.5, 100 mM NaCl, 10 mM $MgCl_2$, 0.1 mM DTT, 5 µg/ml BSA |
| Lb[26] | 25 | 0.324 ± 0.0073 | 0.014 ± 0013 | (-) SC plasmid | 10 mM Tris-HCl, pH 7.5, 100 mM NaCl, 10 mM $MgCl_2$, 0.1 mM DTT, 5 µg/ml BSA |
| Lb* | 30 | 0.199 ± 0.005 | 0.035 ± 0.026 | (-) SC plasmid | 10 mM Tris-HCl, pH 7.5, 50 mM NaCl, 10 mM $MgCl_2$, 5 µg/ml BSA, 0.1 mM DTT |
| As[20] | 25 | 0.052 ± 0.6 | 0.0051 ± 0.2 | Short linear | 50 mM Na-MOPS, pH 7.0, 120 mM NaCl, 5 mM $MgCl_2$, 2 mM DTT |
| As* | 30 | 0.133 ± 0.008 | 0.006 ± 0.002 | (-) SC plasmid | 10 mM Tris-HCl, pH 7.5, 50 mM NaCl, 10 mM $MgCl_2$, 5 µg/ml BSA, 0.1 mM DTT |
| As[25] | 37 | 0.232 ± 0.059 | 0.061 ±0.010 | Short linear | 10 mM Tris-Cl, pH 7.9, 150 mM KCl, 5 mM $MgCl_2$, 1 mM TCEP |
| As[28] | 37 | 0.029 ± 0.002 | 0.11 ± 0.023 | Short linear | 10 mM Tris-HCl, pH 8.0, 100 mM NaCl, 0.9 mM EDTA, 1 mM DTT, 5mM $MgCl_2$ |
| As[28] | 37 | 0.041 ± 0.012 | 0.19 ± 0.011 | Short linear | 10 mM Tris-HCl, pH 8.0, 100 mM NaCl, 0.9 mM EDTA, 1 mM DTT, 10mM $MgCl_2$ |
| As[18] | 37 | 0.10 ± 0.012 | 0.013 ± 0.0012 | Short linear | 50 mM HEPES, pH 7, 150 mM NaCl, 5 mM $MgCl_2$, 2 mM DTT, 0.2 mg/mL BSA |
| Fn* | 30 | 0.561 ± 0.079 | 0.254 ± 0.034 | (-) SC plasmid | 10 mM Tris-HCl, pH 7.5, 50 mM NaCl, 10 mM $MgCl_2$, 5 µg/ml BSA, 0.1 mM DTT |
| Fn[14] | 37 | 0.0075 ± 0.0012 | 0.0011 ± 0.00018 | Short linear | 20 mM Bicine-HCl pH 8, 150mM KCl, 0.5mM TCEP, 5 mM $MgCl_2$ |
| Fn[23] | 37 | 0.0483 ± 0.006 | 0.0338 ± 0.002 | Short linear | 20 mM HEPES, pH 7.5, 150 mM KCl, 5% glycerol, and 0.5 mM DTT, 5 mM $MgCl_2$ |

**Table S1:** *Mean rates of NTS and TS cleavage (with s.d.) from published literature. *Data from this study.*

| | $k_{NTS}$ ($s^{-1}$) | | | $k_{TS}$ ($s^{-1}$) | | |
|---|---|---|---|---|---|---|
| | Mean | S.D. | N | Mean | S.D. | N |
| FnWT | 0.561 | 0.079 | 3 | 0.254 | 0.034 | 3 |
| FnY410A | 0.379 | 0.021 | 3 | 0.988 | 0.089 | 3 |
| FnFLX | 0.362 | 0.048 | 3 | 0.291 | 0.047 | 3 |
| FnΔLoop | 0.072 | 0.008 | 3 | 0.060 | 0.023 | 3 |
| LbWT | 0.199 | 0.005 | 3 | 0.049 | 0.005 | 3 |
| LbW355A | 0.080 | 0.013 | 3 | 0.163 | 0.010 | 3 |
| LbFLX | 0.070 | 0.017 | 3 | 0.114 | 0.031 | 3 |
| LbΔLoop | 0.204 | 0.030 | 3 | 0.066 | 0.007 | 3 |
| AsWT | 0.133 | 0.008 | 3 | 0.006 | 0.002 | 3 |
| AsW382A | 0.092 | 0.012 | 3 | 0.028 | 0.010 | 3 |
| AsFLX | 0.132 | 0.023 | 3 | 0.001 | 1.4e-4 | 3 |
| AsΔLoop | 0.083 | 0.008 | 3 | 0.002 | 8.0e-5 | 3 |
| AsP1153A | 0.103 | 0.024 | 3 | 0.006 | 4.6e-4 | 3 |
| AsFLX-2 | 0.084 | 0.016 | 3 | 0.003 | 0.001 | 3 |

**Table S2:** *Mean rates of NTS and TS cleavage, with standard deviation and n.*

| Cas12a | Amino acids | Mutation |
|---|---|---|
| FnFLX | D1158-D1164 | GSGSGSG substitution (inclusive) |
| FnΔLoop | N1153-T1165 | Deletion (inclusive) |
| LbFLX | K1079-D1085 | GSGSGSG substitution (inclusive) |
| LbΔLoop | R1073-E1088 | Deletion (inclusive) |
| AsFLX | E1165-F1169 | GSGSG substitution (inclusive) |
| AsΔLoop | P1162-Y1173 | Deletion (inclusive) |
| AsP1153A | P1153 | Substitution to alanine |
| AsFLX-2 | E1144-I1155 | Truncation/substitution to GSG |
| **AsΔLoop-2* | *D1137-A1156* | *Deletion (inclusive)* ***no protein expression** |

**Table S3:** *Mutations made to disrupt NUC loops.*

| Orthologue | **Domain** | **Residue #** |
|---|---|---|
| FnCas12a | REC2 | 405-445; 463-469 |
| | NUC loop | 1154-1162 |
| | NUC | 1097-1104; 1196-1203 |
| LbCas12a | REC2 | 345-394 |
| | NUC loop | 1076-1085 |
| | NUC | 1018-1027; 1117-1127 |
| AsCas12a | REC2 | 382-414 |
| | NUC loop | 1169-1172 |
| | NUC | 1088-1096; 1202-1215 |

**Table S4:** *Protein contacts involved in REC2-NUC 'clamping', from MD simulation data.*

| | **Sequence** | **T°(a)** |
|---|---|---|
| **Vector subcloning** | | |
| fn_codingseq_f | AGATATACATATGAGCATCTACCAGGAG | 61 |
| fn_codingseq_r | TCTTCTTTGGGGCATAGTCGGGGACATC | 61 |
| as_codingseq_f | AGATATACATATGACACAGTTCGAGGGC | 64 |
| as_codingseq_r | GGGACATCATTTAGGCATAGTCGGGGAC | 64 |
| pet_fninsert_f | CGACTATGCCCCAAAGAAGAAGCGGAAG | 59 |
| pet_fninsert_r | AGATGCTCATATGTATATCTCCTTCTTAAAGTTAAAC | 59 |
| pet_asinsert_f | CTATGCCTAAATGATGTCCCCGACTATG | 58 |
| pet_asinsert_r | ACTGTGTCATATGTATATCTCCTTCTTAAAGTTAAAC | 58 |
| **Mutagenesis** | | |
| fn_y410a_f | GTTTGATGACGCTTCCGTGATTGGGACC | 66 |
| fn_y410a_r | ACCTGCTGTGACAGGTCT | 66 |
| fn_e1006a_f | GTGGTGTTCGCGGATCTGAAC | 58 |
| fn_e1006a_r | AATGGCATTGTATTCGATG | 58 |
| fn_nucLOOPdel_f | AGGGAGGTGTACCCAACC | 65 |
| fn_nucLOOPdel_r | GATCAGGCGAGATCCGAAG | 65 |
| fn_flexstem_f | CGGGTCCGGGACTAGGGAGGTGTACCCA | 62 |
| fn_flexstem_r | GACCCGGACCCGGAATTTCGAAAGTTGATCAGG | 62 |
| as_w382a_r | AGGGCGCTGCTGATTGTC | 68 |
| as_w382a_f | GTGCGACCACGCGGATACACTGAGGAATG | 68 |
| as_e993a_f | GGTGGTGCTGGCCAACCTGAATTTC | 64 |
| as_e993a_r | ACGGCCTGGTAGTGGATC | 64 |
| as_p1153a_f | CAAGGGCACCGCGTTCATCGCCG | 67 |
| as_p1153_r | GCGTCAAACTGTGTCTCGTTC | 67 |
| as_flx2_f | TGGCGCCGGCAAGAGAATCGTG | 65 |
| as_flx2_r | GAGCCGTTCTTCTCGAACACGATATCCC | 65 |

| | | |
|---|---|---|
| as_nucLOOPdel_f | CGGGACCTGTATCCTGCC | 67 |
| as_nucLOOPdel_r | CACGATTCTCTTGCCGGC | 67 |
| as_flexstem_f | CGGGTCCACCGGCAGATACCGGGAC | 69 |
| as_flexstem_r | GACCCGGAGATCACTGGCACGATTCTCTTGC | 69 |
| lb_w355a_f | CTTCGGCGAGGCGAACGTGATC | 56 |
| lb_w355a_r | ATATCCTTGGAGATTGTG | 56 |
| lb_e925a_f | ATCGCCCTGGCGGACCTGAAC | 64 |
| lb_e925a_r | CACGGCATCGTACTTCTCCAC | 64 |
| lb_nucLOOPdel_f | GTGTGCCTGACCAGCGC | 69 |
| lb_nucLOOPdel_r | GATCCGGTTGCCGTAGGAG | 69 |
| lb_flxstem_f | CGGGTCCGGGTGGGAGGAGGTGTGCCTG | 68 |
| lb_flxstem_r | GACCCGGACCCAGGATTCCGGAAGATTCTGATCCGG | 68 |

**Table S5:** Sequences of cloning primers.

| Name | Sequence |
|---|---|
| Mini_locus_top | ATAAGGAGATATACCATGGGAATTTCTACTGTTGTAGATTATGGGTATAAATGGGCTCGCGAAATTTCTACTGTTGTAGATTATGGGTATAAATGGGCTCGCGAAATTTCTACTGTTGTAGATGAATTCGAGCTCGGCGCGCC |
| Mini_locus_bottom | GGCGCGCCGAGCTCGAATTCATCTACAACAGTAGAAATTTCGCGAGCCCATTTATACCCATAATCTACAACAGTAGAAATTTCGCGAGCCCATTTATACCCATAATCTACAACAGTAGAAATTCCCATGGTATATCTCCTTAT |
| FQ ssDNA | 5'-/56-FAM/TTT TTT TTT/ZEN/TTT/3IaBkFQ/ -3' |
| TS | CCCGGTGTCACGCCACTTGACAGGCGAGTAACAGACATGGACCATCAGGAAACATTAACGTACTGATGTTAACAGCTGACCCAATAAGTGGCAGAG |
| TS_truncated | TAACAGACATGGACCATCAGGAAACATTAACGTACTGATGTTAACAGCTGACCCAATAAGTGGCAGAG |

**Table S6:** Oligonucleotide sequences used in 'locus' cloning and *trans* cleavage assays.

| **crRNAs for in vitro cis and trans cleavage assays** | |
|---|---|
| Lb_23 | UAAUUUCUACUAAGUGUAGAUCUGAUGGUCCAUGUCUGUUACUC |
| Lb_20 | UAAUUUCUACUAAGUGUAGAUCUGAUGGUCCAUGUCUGUUA |
| As_23 | UAAUUUCUACUCUUGUAGAUCUGAUGGUCCAUGUCUGUUACUC |
| As_20 | UAAUUUCUACUCUUGUAGAUCUGAUGGUCCAUGUCUGUUA |
| Fn_23 | UAAUUUCUACUGUUGUAGAUCUGAUGGUCCAUGUCUGUUACUC |

| | |
|---|---|
| Fn_20 | UAAUUUCUACUGUUGUAGAUCUGAUGGUCCAUGUCUGUUA |
| **crRNAs for human cell gene editing via RNP electroporation** | |
| Lb_DNMT1-3 | UAAUUUCUACUAAGUGUAGAUCUGAUGGUCCAUGUCUGUUACUCG |
| Lb_DNMT1-7 | UAAUUUCUACUAAGUGUAGAUGCUCAGCAGGCACCUGCCUCAGCU |
| Lb_AGBL1 | UAAUUUCUACUAAGUGUAGAUGAUUGAAGGAAAAGUUACAAAGGU |
| As_DNMT1-3 | UAAUUUCUACUCUUGUAGAUCUGAUGGUCCAUGUCUGUUACUCG |
| As_DNMT1-7 | UAAUUUCUACUCUUGUAGAUGCUCAGCAGGCACCUGCCUCAGCU |
| As_AGBL1 | UAAUUUCUACUCUUGUAGAUGAUUGAAGGAAAAGUUACAAAGGU |
| Fn_DNMT1-3 | UAAUUUCUACUGUUGUAGAUCUGAUGGUCCAUGUCUGUUACUCG |
| Fn_DNMT1-7 | UAAUUUCUACUGUUGUAGAUGCUCAGCAGGCACCUGCCUCAGCU |
| Fn_AGBL1 | UAAUUUCUACUGUUGUAGAUGAUUGAAGGAAAAGUUACAAAGGU |

**Table S7:** RNA oligonucleotide sequences.

| Site name | site sequence | primer set name* | Forward primer | Reverse primer |
|---|---|---|---|---|
| dnmt1-3 | TTTCCTGATGGTCCATGTCTGTTA | DNMT1-3 / TTTC-2 | TCCCTTAGCACTCTGCCACTTAT | GTTAAAAACACAACATCAGTGCATGT |
| dnmt1-7 | TTTGGCTCAGCAGGCACCTGCCTC | DNMT1-7 /TTTG-2 | AGCAGGCCTTTGGTCAGGTT | AGACATGGACCATCAGGAAACAT |
| agbl1 | TTTAGATTGAAGGAAAAGTTACAA | AGBL1 /TTTA-6 | GAAGAGAAATCTGCGTGGAGAGA | GAAAACCCCCAAAAATCCCA |
| dnmt1-3-OT1 | TTTCCTGATGGTCCATGTCTGAAT | WT-AsCas12a-TTTC-2-chr1 | CCTATTCTTCCGCCATTTTCC | CTGAATGCCTGCTATGTACACACA |
| dnmt1-3- | TTTCCTGATGGTCCACATCTGTTA | enAsCas12a-TTTC- | TGCCACATTACCCTCTAAAAGTCA | TGCGCAGTGTGTTTAGGAAGTT |

| | | | | |
|---|---|---|---|---|
| OT 2 | | 2-chr11 | | |
| dnmt1-3-OT 3 | TTTCCTGATGGTCCATATCTGTGG | enAsCas12a-TTTC-2-chr10 | TCAGGTGATGGTTTGGCAATC | ATAGAAAGCCTCCCCACCTAAGG |
| dnmt1-3-OT 4 | TTTCCTGATGGTCCATACCTGTTA | enAsCas12a-TTTC-2-chrX | CTCTTCCCCTCAACCACTAAATATG | TCTGAGGTTTTCTTCCCTTTTCC |
| dnmt1-7-OT 1 | AGCAGCTCAGCAGGCACCTGCCTT | enAsCas12a-TTTG-2-chr21 | CAGCCAGGGCCTCATTAAAC | CATGTTGAATGTTGGCACGAA |
| dnmt1-7-OT 2 | AACAGCTCAGCAGACACCTGCCAA | enAsCas12a-TTTG-2-chr2 | TTGCCATGACAACAACTCAGG | GGACGCAATGTCAGCTGTGT |
| dnmt1-7-OT 3 | TTTAGCTCAGCTGACACCTGCCCA | enAsCas12a-TTTG-2-chr12 | TGTTGCATAGGCTTGTTCTGTTC | TTGTTAGGGTTCATTGTAGCTGACA |
| abgl1-OT 1 | TTTAGATTAATGGAAAAGTTACAA | enAsCas12a-TTTA-6-chr6 | CACTGCACCCGGCCATA | TGTGGAGTCTTGAACTGGATCTTG |
| abgl1-OT 2 | TTTAGACTTAAAGAAAAGTTACAA | enAsCas12a-TTTA-6-chr8 | GTGGTAGTACTTCCTTCATACTGAGCA | GTCCCGTGGAGTAGTTGATGTG |
| abgl1-OT 3 | CTTGGATTAAAGGAAAAGCTACAA | enAsCas12a-TTTA-6-chr10 | TGGAAATAAGCAATCAGGCCTT | TTTGTTTTGTCTTTCCTTACATCGTT |

**Table S8:** Primers for high-throughput sequencing. *Sites names as per Kleinstiver 2019

"Kleinstiver, B.P., Sousa, A.A., Walton, R.T. et al. Engineered CRISPR–Cas12a variants with increased activities and improved targeting ranges for gene, epigenetic and base editing. Nat Biotechnol 37, 276–282 (2019). https://doi.org/10.1038/s41587-018-0011-0"
